# Protect TUDCA stimulated CKD-derived hMSCs against the CKD-Ischemic disease via upregulation of PrP^C^

**DOI:** 10.1101/401356

**Authors:** Yeo Min Yoon, SangMin Kim, Yong-Seok Han, Chul Won Yun, Jun Hee Lee, Hyunjin Noh, Sang Hun Lee

## Abstract

Although autologous human mesenchymal stem cells (hMSCs) are a promising source for regenerative stem cell therapy, the barriers associated with pathophysiological conditions in this disease limit therapeutic applicability to patients. We proved treatment of CKD-hMSCs with TUDCA enhanced the mitochondrial function of these cells and increased complex I & IV enzymatic activity, increasing PINK1 expression and decreasing mitochondrial O_2_^•−^ and mitochondrial fusion in a PrP^C^-dependent pathway. Moreover, TH-1 cells enhanced viability when co-cultured *in vitro* with TUDCA-treated CKD-hMSC. *In vivo*, tail vein injection of TUDCA-treated CKD-hMSCs into the mouse model of CKD associated with hindlimb ischemia enhanced kidney recovery, the blood perfusion ratio, vessel formation, and prevented limb loss, and foot necrosis along with restored expression of PrP^C^ in the blood serum of the mice. These data suggest that TUDCA-treated CKD-hMSCs are a promising new autologous stem cell therapeutic intervention that dually treats cardiovascular problems and CKD in patients.

## Introduction

Despite the advancement of therapeutic research, chronic kidney disease (CKD) remains a huge public health burden, afflicting approximately 7.7% of the US population.(Castro & Coresh, 2009) Patients with CKD fail to excrete toxic metabolites and organic waste solutes that are normally removed by healthy kidneys, with the resultant accumulation of such toxins in these patients and significantly increased risks of other diseases including anemia, metabolic bone disease, neuropathy, and cardiovascular disease.(D’Hooge et al, 2003; Gabriele et al, 2016; Meijers et al, 2010) Vascular diseases (VDs) remain a major problem frequently associated with CKD, and mortality among individuals with CKD is attributable more to secondary vascular damage than to actual kidney failure.(Sarnak et al, 2003a) VDs in patients with CKD are traditionally known to be caused by increased risks of arterial hypertension due to aberrations in CKD, but recent research shows evidence of novel risk factors such as premature senescence, decreased proliferation, and apoptosis of cardiovascular cell types in relevant tissues owing to oxidative stress induced by uremic toxins circulating in the serum of patients with CKD.(Schlieper et al, 2016) Despite the overbearing burden, currently available therapies for CKD often lead to adverse side effects and malignant pharmacokinetic changes.(Velenosi & Urquhart, 2014)

Mesenchymal stem cells (MSCs), a subcategory of adult stem cells, possess excellent therapeutic potentials because of their ability to secrete reparative factors and cytokines, their capacity for tissue-specific differentiation, and their self-renewal ability, especially apt for organ regeneration and tissue repair.(Rogers et al, 2016; Uccelli et al, 2008) In particular, some studies have delineated the benefits of MSC therapy as a promising modality targeting VD associated with CKD.(Lee et al, 2016; Little, 2006) The paracrine effect of MSCs, include cell migration, homing, and stimulation, and antiapoptosis, and antioxidantive factor for repair damaged tissue, as well as the therapeutic potentials of MSCs related to neovascularization, making MSCs a promising cell source with strong regenerative as well as modulatory abilities for alleviation of both renal and vascular complications.(Lee et al, 2016; Liang et al, 2014; Rogers et al, 2016; Uccelli et al, 2008) Nonetheless, one of the major drawbacks of MSC therapy is that MSCs obtained from patients with CKD are exposed to endogenous uremic toxins that decrease their viability and therapeutic potential.(Klinkhammer et al, 2014)

Tauroursodeoxycholic acid (TDUCA), a conjugate of ursodeoxycholic acid (UDCA) and taurine, is a bile acid produced in humans in small amounts and is approved by the US Food and Drug Administration to be safely used as a drug for human intake.(Yoon et al, 2016) Cellular prion protein (PrP^C^), a glycoprotein commonly known to pathogenically misfold into PrP^SC^ (which causes neurodegenerative disorders(Prusiner, 1982; Prusiner et al, 1998)) plays vital metabolic roles in the body such as regulation of cell differentiation, cell migration, and long-term potentiation of synapses.(Besnier et al, 2015; Hornshaw et al, 1995; Pinheiro et al, 2015) Furthermore, PrP^C^ possesses a therapeutic function: it increases survival of MSCs, and one study has shown that PrP^C^ is essential for reducing the oxidative stress at the injury site of an ischemia model.(Yoon et al, 2016) More specifically, some studies have suggested that PrP^C^ may play a role in mitochondrial functions involving levels of oxidative stress and mitochondrial morphology,(Lobao-Soares et al, 2005; Miele et al, 2002) and TUDCA has been shown to be neuroprotective in a model of acute ischemic stroke via stabilization of the mitochondrial membrane.(Soares et al, 2018; Sola et al, 2002) Nevertheless, the precise mechanism by which TUDCA potentiates MSCs via PrP^C^-dependent enhancement of mitochondrial function is not yet investigated.’

PINK1 is well known recruit Parkin to depolarized mitochondria, then to increase activation of Parkin E3 ligase activity, resulting in ubiquitylation of multiple substrates at the outer mitochondrial membrane. It takes a role in critical upstream step mitophagy, destruction of damage mitochondria.(Gladkova et al, 2018; Lazarou et al, 2012) PINK1 also regulated mitochondrial respiration driven by electron transport chain (ETC) complex ? & IV activity.(Liu et al, 2011) Thus, PINK1 is a key factor for retain mitochondria potential.

In our study, we aimed to elucidate the therapeutic benefits of TUDCA-treated CKD- hMSCs for targeting VD associated with CKD. We investigated the precise mechanism via which TUDCA enhanced the survival of CKD-hMSCs in CKD-associated VD: through mitochondrial membrane potential enhancement via the PrP^C^–PINK1 axis. We hypothesized that PrP^C^ is a protein of interest for the treatment of patients with VD associated with CKD, and that PrP^C^-mediated mitochondrial recovery from oxidative stress (common in patients with CKD) is a crucial factor for therapeutic targeting of CKD-related VD.

## Results

### Patients with CKD have lower levels of serum PrP^C^, and CKD patient–derived MSCs show decreased mitochondrial membrane potential

To analyze biomarkers of CKD, we measured estimated glomerular filtration rate in the serum of the patients with CKD (n = 25, Healthy control; n = 10). We confirmed that our patients with CKD were at stage 3b–5 (average eGFR: 22.4; Fig 1A). In patients with CKD, ELISA results revealed a decreased level of PrP^C^ in serum (Fig 1B). Patients with CKD showed a decreased slope on the eGFR axis against concentration on PrP^C^ axis as compared to the healthy control group (Fig 1C). Thus, we investigated whether the reduced expression of PrP^C^ may be related to CKD. It has been known that mitochondrial PrP^C^ maintains mitochondrial membrane potential and reduces accumulation of ROS to retain activity of the electron transport chain complex.(Brown et al, 2002; Lobao-Soares et al, 2005; Stella et al, 2012) To confirm obtained MSCs from healthy and patient group sustained similar normally MSCs, we progressed differentiate into chondrogenic, adipogenic, and osteogenic cells, and measured surface marker CD11b, and CD45-negative control, and CD44, and Sca-1, positive control (Fig EV1). In contrast to Healthy-hMSCs, CKD-hMSCs (n = 4) showed decreased expression of PrP^C^ by using ELISA (Fig 1D). CKD-hMSCs (n = 4) also had weaker binding of PrP^C^ to PINK1 (Fig 1E and F). These results indicated that CKD-hMSCs might mediate mitochondrial dysfunction by decreased binding of PrP^C^ and PINK1. To confirm that mitochondrial dysfunction in CKD-hMSCs (n = 3) is caused by decreased mitochondrial PrP^C^ levels, we analyzed accumulated ROS in mitochondria and complex I & IV activities by ELISAs and FACS. CKD-hMSCs accumulated more O_2_^•−^ (superoxide) in mitochondria according to the MitoSOX assay and showed decreased complex ? & IV activities (Fig 1G–I).

**Figure 1.**
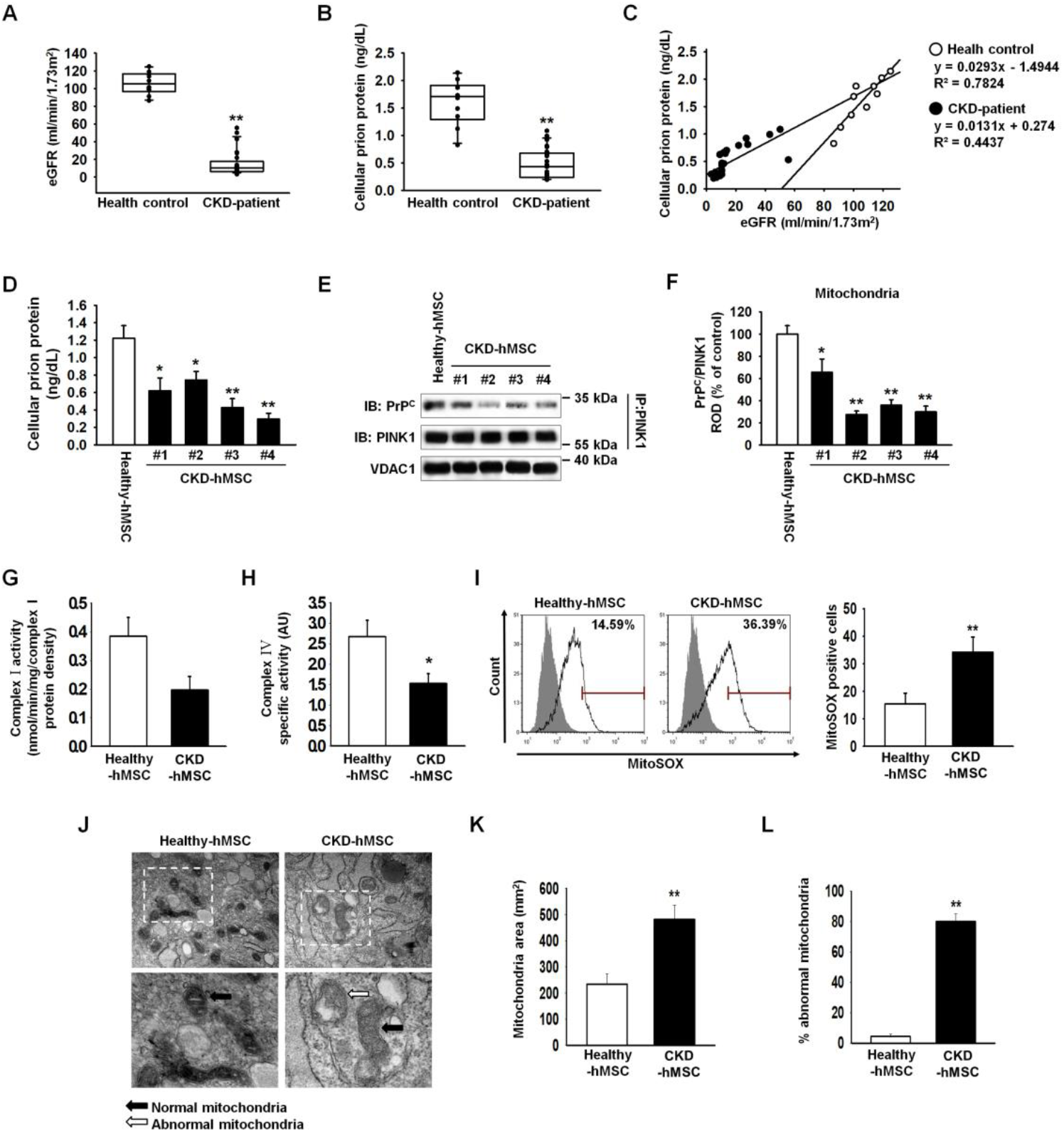
Patients with CKD show a decreased level of PrP^C^, and CKD patient–derived MSCs show decreased mitochondrial functionality. **A, B.** Patients with CKD and healthy control groups (Healthy control): estimated glomerular filtration rate (eGFR), and PrP^C^ in each group in their serum (100 μL samples). Values represent the mean ± SEM. \**p* < 0.05, and \*\**p* < 0.01 vs. Healthy control. **C.** A graph on eGFR-axis against the concentration on the PrP^C^-axis. **D.** A Healthy-hMSCs and CKD-hMSCs measured concentrations of PrP^C^ by ELISA. Values represent the mean ± SEM. \**p* < 0.05, and \*\**p* < 0.01 vs. untreated Healthy-hMSCs **E.** Immunoprecipitates with an anti-PINK1 antibody were analyzed for Healthy-hMSCs and CKD-hMSCs by western blotting using an antibody that recognized PrP^C^. **F.** The level of PrP^C^, whose binding with PINK1 was normalized to that of VDAC1. Values represent the mean ± SEM. \*\**p* < 0.01 vs. Healthy-hMSCs. **G**, **H.** Complex ? and complex ? activities were analyzed in Healthy-hMSCs and CKD- hMSCs by ELISAs. Values represent the mean ± SEM. \**p* < 0.05 vs. Healthy-hMSCs. **I.** The positive signals of MitoSOX were quantified by FACS analysis with staining of Healthy-hMSCs and CKD-hMSCs. Values represent the mean ± SEM. \*\**p* < 0.01 vs. Healthy-hMSCs. **J.** Representative TEM images (*n* = 3 per group) of Healthy-hMSCs and CKD-hMSCs. Scale bars are shown enlarged on the left. Scale bars: 500 nm. **K.** Quantitative analyses of morphometric data from TEM images. Values represent the mean ± SEM. \*\**p* < 0.01 vs. Healthy-hMSCs. **L.** Percentages of abnormal mitochondria that were swollen with evidence of severely disrupted cristae throughout a mitochondrion) obtained from a TEM image. Values represent the mean ± SEM. \*\**p* < 0.01 vs. Healthy-hMSCs.

We also found that CKD-hMSCs contained increased numbers of abnormal mitochondria and increased mitochondrial area, which indicated greater mitofusion, according to transmission electron microscopy (TEM; Fig 1J–L). To confirm that mitochondrial dysfunction was dependent on mitochondrial PrP^C^, we demonstrated that a knockdown of PrP^C^ in Healthy- hMSCs increased mitochondrial O_2_^•−^ (superoxide) amounts and decreased complex I & IV activities because of decreased PrP^C^ binding to PINK1 (Fig EV2A-D). Our results certainly indicated that in patients with CKD, decreased PrP^C^ amounts influenced mitochondrial PrP^C^, which normally might maintain mitochondrial membrane potential through binding to PINK1.

### TUDCA enhances mitochondrial membrane potential via increasing the level of mitochondrial PrP^C^ in CKD-hMSCs

We verified that patients with CKD underexpress PrP^C^. This phenomenon suggests that CKD-hMSCs may harbor decreased mitochondrial membrane potential and increased numbers of abnormal mitochondria via a PrP^C^-dependent mechanism. Thus, increasing the expression of PrP^C^ may effectively influence not only mitochondrial membrane potential but also stem cell viability. In our previous study, we verified that TUDCA-treated MSCs are protected against ROS-induced cell apoptosis via increased Akt-PrP^C^ signaling.(Yoon et al, 2016) Nevertheless, we do not know whether TUDCA affects mitochondrial membrane potential through increased expression of PrP^C^. To verify our theory, i.e., to increase PrP^C^ expression for enhancing mitochondrial membrane potential and stem cell therapy through treatment with TUDCA, Healthy-hMSCs or CKD-hMSCs were treated with TUDCA and showed an increased level of PrP^C^ as compared with no treatment with TUDCA. TUDCA- treated CKD-hMSCs, particularly, showed a similar levels relative to Healthy-hMSCs (Fig 2A). However, after pretreatment Heathy-hMSCs or CKD-hMSCs with Akt inhibitor, disappeared expression PrP^C^ protein-induced TUDCA. These results indicated that our theory is correct: stem cell therapy with TUDCA-treated CKD-hMSCs should be as effective as stem cell therapy with Healthy-hMSCs, therefore TUDCA regulated expression of PrP^C^ via Akt signal patyway. We showed that TUDCA-treated Healthy-hMSCs or CKD-hMSCs harbor similarly increased levels of catalase and SOD activities, which are antioxidative enzymes (Fig EV3A and B). These results suggested that TUDCA increased the expression of normal cellular prion protein. To confirm that PrP^C^ increased mitochondrial membrane potential via treatment with TUDCA, Healthy-hMSCs or CKD-hMSCs were treated with TUDCA, and we observed increased expression of PrP^C^ in mitochondria (Fig 2B and 2C). To verify mitochondrial membrane potential and mitophagy, we measured expression of PINK1, which regulates mitochondrial membrane potential by increased complex I & IV activities and is located in the outer mitochondrial membrane for initiation of mitophagy of dysfunctional mitochondria.(Narendra & Youle, 2011) In contrast to TUDCA-treated Healthy-hMSCs, TUDCA-treated CKD-hMSCs showed significantly increased expression of PINK1 (Fig 2B and C). TUDCA-treated Healthy-hMSCs or CKD-hMSCs manifested higher complex I & IV activities as compared to the group without TUDCA treatment (Fig EV3C and D). These results indicated that the key mechanism of TUDCA effectiveness for mitochondrial membrane potential is mitochondrial PrP^C^ (rather than PINK1). To confirm that TUDCA-treated CKD-hMSCs had better mitochondrial membrane potential via the binding of mitochondrial PrP^C^ to PINK1, we analyzed TUDCA-treated CKD-hMSCs by immunoprecipitation experiments (Fig 2D and E). TUDCA-treated CKD-hMSCs also showed increased complex I & IV activities and decreased mitochondrial O_2_^•−^ amounts (Fig 2F–H). In contrast, the knockdown of PrP^C^ in TUDCA-treated CKD-hMSCs increased mitochondrial O_2_^•−^ levels and decreased complex I & IV activities because of decreased binding of PrP^C^ and PINK1 (Fig 2E–H).

**Figure 2.**
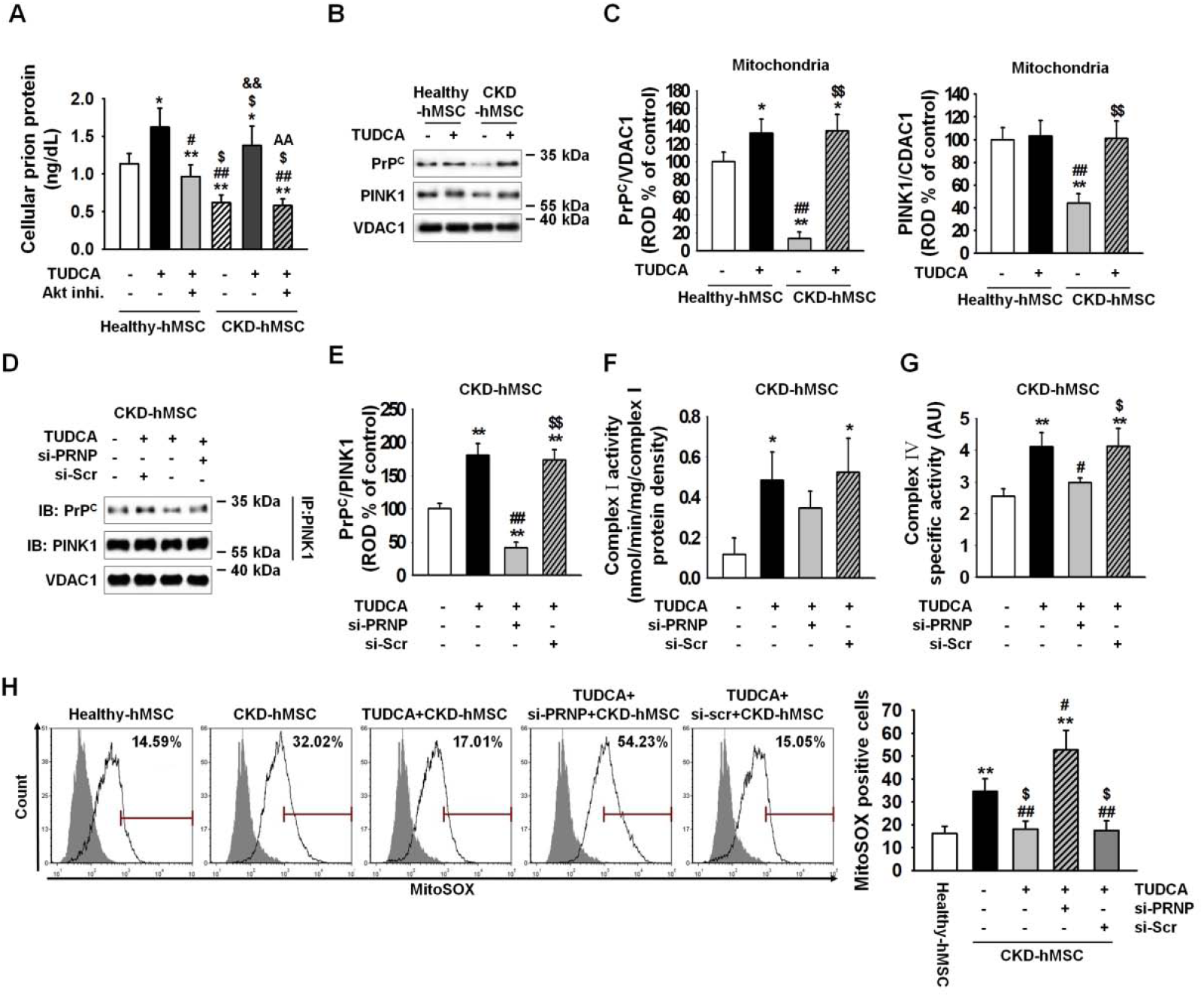
TUDCA enhanced mitochondrial membrane potential through increased mitochondrial PrP^C^ levels in CKD-hMSCs. **A.** Healthy-hMSCs and CKD-hMSCs were treated with TUDCA, and we measured concentrations of PrP^C^ by ELISA. Values represent the mean ± SEM. \*\**p* < 0.01 vs. untreated Healthy-hMSCs, ##*p* < 0.01 vs. TUDCA treatment of Healthy-hMSCs, $$*p* < 0.01 vs. untreated CKD-hMSCs. **B.** Western blot analysis quantified the expression of PrP^C^ and PINK1 after treatment with TUDCA of Healthy-hMSCs and CKD-hMSCs. **C.** The expression levels were determined by densitometry relative to VACD1. Values represent the mean ± SEM. \**p* < 0.05 and \*\**p* < 0.01 vs. untreated Healthy-hMSCs, ##*p* < 0.01 vs. TUDCA treatment of Healthy-hMSCs, $$*p* < 0.01 vs. untreated CKD-hMSCs. **D.** Immunoprecipitates with anti-PINK1 were analyzed after treatment of TUDCA-pretreated CKD-hMSCs with si-PRNP by western blot using an antibody that recognized PrP^C^. **E.** The level of PrP^C^, whose binding with PINK1 was normalized to that of VDAC1. Values represent the mean ± SEM. \*\**p* < 0.01 vs. untreated CKD-hMSCs, ##*p* < 0.01 vs. TUDCA treatment of CKD-hMSCs, $$*p* < 0.01 vs. TUDCA-treated CKD-hMSCs pretreated with si- PRNP. **F, G.** Complex ? and complex IV activities were analyzed by an ELISA after treatment of TUDCA-pretreated CKD-hMSCs with si-PRNP. Values represent the mean ± SEM. \**p* < 0.05, and \*\**p* < 0.01 vs. Healthy-hMSCs, #*p* < 0.05 vs. TUDCA treatment of CKD-hMSCs, $*p* < 0.05 vs. TUDCA-treated CKD-hMSCs pretreated with si-PRNP. **H.** The positive signals of MitoSOX were quantified by FACS analysis with staining after treatment of TUDCA-pretreated CKD-hMSCs with si-PRNP. \*\**p* < 0.01 vs. Healthy-hMSCs, #*p* < 0.05, and ##*p* < 0.01 vs. TUDCA treatment of CKD-hMSCs, $*p* < 0.05 vs. TUDCA- treated CKD-hMSCs pretreated with si-PRNP.

### TUDCA protects CKD-hMSCs against dysfunctional mitochondria via increased mitophagy and increased cell proliferation

To clarify whether the reduced mitochondrial membrane potential was caused by a decrease in binding of mitochondrial PrP^C^ to PINK1, we measured complex I & IV activities and mitochondrial O_2_^•−^ concentrations. These results indicated that TUDCA-treated CKD-hMSCs had fewer dysfunctional mitochondria. Mitophagy decreased the number of dysfunctional mitochondria, whereas mitofussion may be increased in abnormal conditions, such as CKD, inflammation, and oxidative stress, and reduces mitophagy signaling.(Schlieper et al, 2016) We confirmed that CKD-hMSCs contained a larger mitochondria area on cross-sections as well as abnormal mitochondria, whereas TUDCA treatment decreased the mitochondrial area and the number of abnormal mitochondria in CKD hMSCs according to TEM (Fig 3A–C). These results indicated that the effect of TUDCA reduced dysfunctional mitochondria numbers by decreasing mitofusion and increasing mitophagy. We verified downregulation of mitofission suppressor protein p-DPR1 (phosphorylation Ser 637) and of mitofusion- associated protein MFN1, and OPA1 (regulated by p-DRP1) in TUDCA-treated CKD- hMSCs by western blot analysis (Fig 3D and E). TUDCA-treated CKD-hMSCs also showed increased mitophagy because we analyzed autophagy-regulated proteins, P62 and LC3B, by western blotting of the mitochondrial fraction (Fig 3D and E). In contrast, knockdown PrP^C^ CKD-hMSCs did not respond to TUDCA and did not show enhancement of mitochondrial function according to TEM (Fig 3A–C). TUDCA-treated CKD-hMSCs after a knockdown of PrP^C^, revealed greater activation of p-DPR1 and expression of MFN1 and OPA1 (Fig 3D and 3E), and decreased mitophagy, increased expression P62, and decreased expression of LC3B II (Fig 3f and 3g). These results supported our argument that the knockdown of PrP^C^ in Healthy-hMSCs increased the mitochondrial area and numbers of abnormal mitochondria. These results indicated that TUDCA treatment of CKD-hMSCs reduced mitochondrial fission and increased mitophagy to decrease dysfunction of mitochondria. These data confirmed that TUDCA protects mitochondrial membrane potential and reduces dysfunction of mitochondria through activation of mitophagy. To analyze the increase in cell proliferation by TUDCA treatment of Healthy-hMSCs and CKD-hMSCs, we conducted a BrdU assay, cell cycle assay, and evaluation of CDK4–Cyclin D1 activity (Fig 4A–C). The knockdown of PrP^C^ indicated that the effects of TUDCA downregulated cell cycle–associated proteins, CDK2, cyclin E, CDK4, and cyclin D1, according to western blotting (Fig 4D and E). To analyze the attenuation of cell proliferation by the dysfunction of mitochondria, we also conducted a BrdU assay and found reduced cellular proliferation as well as the decreased S phase activity according to FACS, and CDK4/cyclin D1 activity (Fig 4F–H). These results showed that CKD-hMSCs slowed down the proliferation due to deficiency of mitochondrial function under the influence of the PrP^C^ knockdown and increased accumulation of mitochondrial O_2_^•−^ and decreased complex I & IV activities.

**Figure 3.**
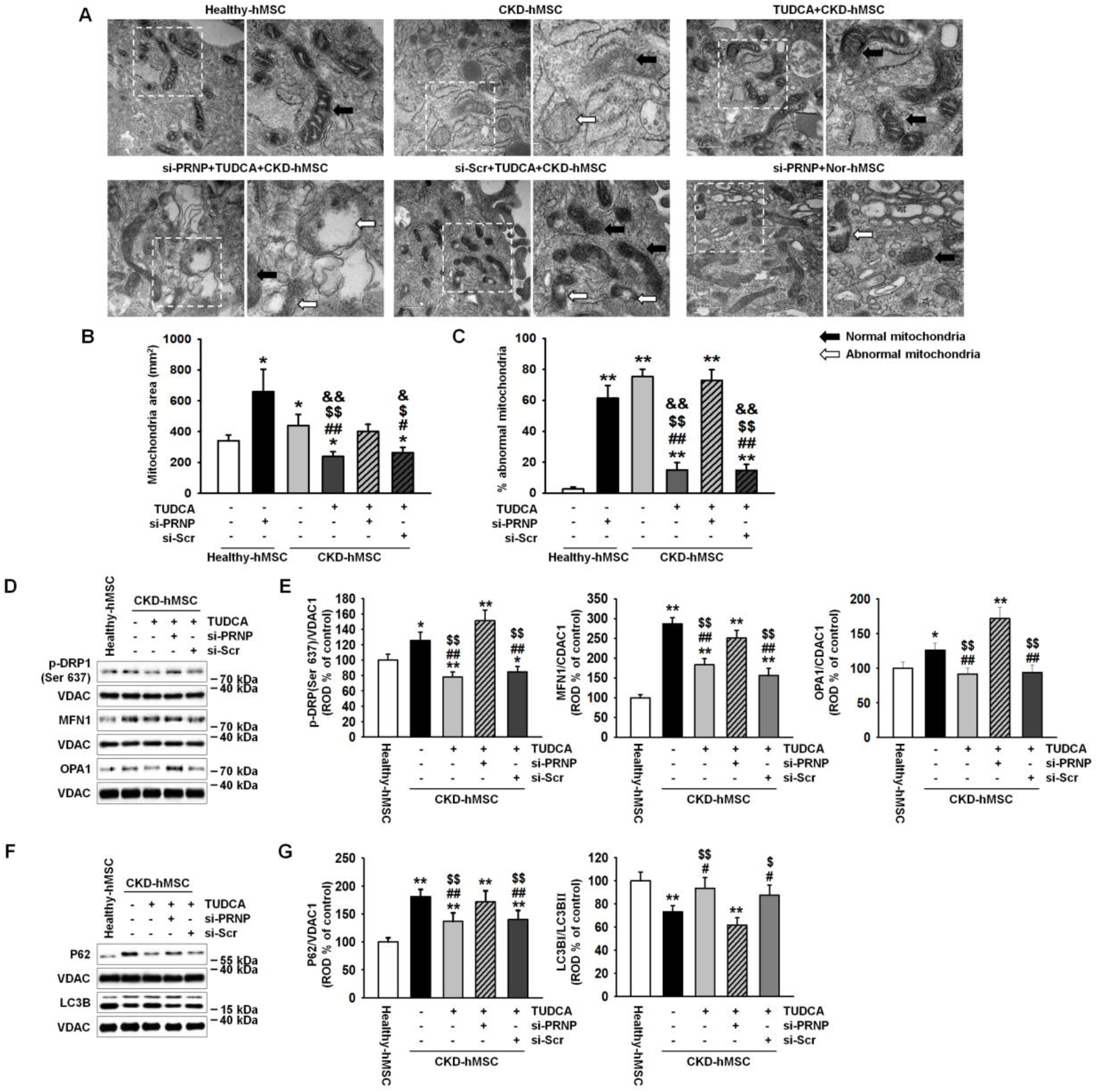
TUDCA regulated mitochondrial fission via increased mitochondrial PrP^C^ amounts in CKD-hMSCs. **A.** Representative TEM images (*n* = 3 per group) after incubation of Healthy-hMSCs with or without si-PRNP or after treatment of TUDCA-pretreated CKD-hMSCs with si-PRNP. Scale bars are shown enlarged on the left. Scale bars: 500 nm. **B.** Quantitative analyses of morphometric data from TEM images. Values represent the mean ± SEM. \**p* < 0.05 vs. untreated Healthy-hMSCs, #*p* < 0.05 and ##*p* < 0.01 vs. si-PRNP-treated Healthy-hMSCs, $*p* < 0.05 and $$*p* < 0.01 vs. CKD-hMSCs, &*p* < 0.05 and &&*p* < 0.01 vs. TUDCA-treated CKD-hMSCs pretreated with si-PRNP. **C.** Percentages of abnormal mitochondria, which were swollen with evidence of severely disrupted cristae throughout a mitochondrion) obtained from TEM images. Values represent the mean ± SEM. \*\**p* < 0.01 vs. untreated Healthy-hMSCs, ##*p* < 0.01 vs. si-PRNP-treated Healthy-hMSCs, $$*p* < 0.01 vs. CKD-hMSCs, &&*p* < 0.01 vs. TUDCA-treated CKD-hMSCs pretreated with si-PRNP. **D.** Western blot analysis quantified the expression of p-DRP1, MFN1, and OPA1 in Healthy- hMSCs, treatment of CKD-hMSCs with or without TUDCA, and after treatment of TUDCA- pretreated CKD-hMSCs with si-PRNP. **E.** The expression levels were determined by densitometry relative of VACD1. Values represent the mean ± SEM. \**p* < 0.05 and \*\**p* < 0.01 vs. Healthy-hMSCs, #*p* < 0.05 and ##*p* < 0.01 vs. CKD-hMSCs, $$*p* < 0.01 vs. TUDCA-treated CKD-hMSCs pretreated with si- PRNP. **F.** Western blot analysis quantified the expression of P62, and LC3B in Healthy-hMSCs, treatment CKD-hMSCs with or without TUDCA, and after treatment of TUDCA-pretreated CKD-hMSCs with si-PRNP. **G.** The expression levels were determined by densitometry relative to VACD1. Values represent the mean ± SEM. \*\**p* < 0.01 vs. Healthy-hMSCs, #*p* < 0.05 and ##*p* < 0.01 vs. CKD-hMSCs, $*p* < 0.05 and $$*p* < 0.01 vs. TUDCA-treated CKD-hMSCs pretreated with si- PRNP.

**Figure 4.**
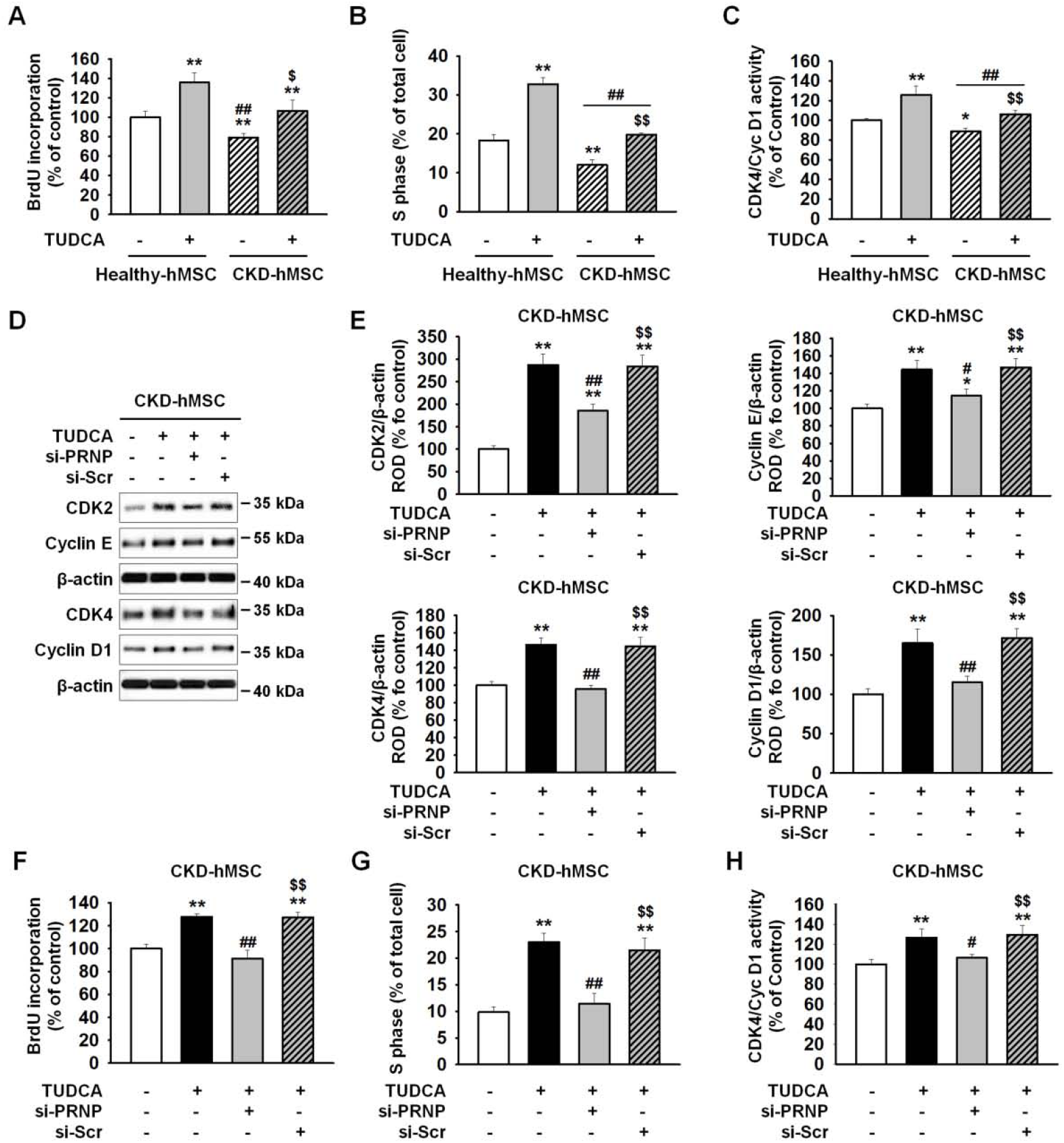
TUDCA treatment of CKD-hMSCs enhanced cell proliferation through upregulation of PrP^C^. **A.** Cell proliferation of Healthy-hMSCs or CKD-hMSCs analyzed after treatment with or without TUDCA. Values represent the mean ± SEM. \*\**p* < 0.01 vs. untreated Healthy-hMSCs, ##*p* < 0.01 vs. TUDCA-treated Healthy-hMSCs, $*p* < 0.05 vs. untreated CKD-hMSCs. **B.** The number of S phase Healthy-hMSCs or CKD-hMSCs was determined by FACS analysis of PI-stained cells. Values represent the mean ± SEM. \*\**p* < 0.01 vs. untreated Healthy-hMSC, ##*p* < 0.01 vs. TUDCA-treated Healthy-hMSCs, $$*p* < 0.01 vs. untreated CKD-hMSCs. **C.** CKD4/Cyclin D1 activity was analyzed by ELISA. Values represent the mean ± SEM. \**p* < 0.05, and \*\**p* < 0.01 vs. untreated Healthy-hMSCs, ##*p* < 0.01 vs. TUDCA-treated Healthy-hMSCs, $$*p* < 0.01 vs. untreated CKD-hMSCs. **D.** Western blot analysis quantified the cell cycle–associated proteins, CDK2, Cyclin E, CDK4, and cyclin D1 after treatment of CKD-hMSCs with or without TUDCA, and after treatment of TUDCA-pretreated CKD-hMSCs with si-PRNP. **E.** The expression levels were determined by densitometry relative of β-actin. Values represent the mean ± SEM. \**p* < 0.05 and \*\**p* < 0.01 vs. CKD-hMSCs, #*p* < 0.05 and ##*p* < 0.01 vs. TUDCA-treated CKD-hMSCs, $$*p* < 0.01 vs. TUDCA-treated CKD-hMSCs pretreated with si-PRNP. **F.** Cell proliferation of CKD-hMSCs analyzed after treatment of TUDCA-pretreated CKD- hMSCs with si-PRNP by a BrdU assay. Values represent the mean ± SEM. \*\**p* < 0.01 vs. untreated CKD-hMSCs, ##*p* < 0.01 vs. TUDCA-treated CKD-hMSCs, $$*p* < 0.01 vs. TUDCA-treated CKD-hMSCs pretreated with si-PRNP. **G.** The number of S phase CKD-hMSCs was determined by FACS analysis of PI-stained cells. Values represent the mean ± SEM. \*\**p* < 0.01 vs. untreated CKD-hMSCs, ##*p* < 0.01 vs. TUDCA-treated CKD-hMSCs, $$*p* < 0.01 vs. TUDCA-treated CKD-hMSCs pretreated with si-PRNP. **H.** CKD4/Cyclin D1 activity was analyzed by ELISA. Values represent the mean ± SEM. \*\**p* < 0.01 vs. untreated CKD-hMSCs, #*p* < 0.05 vs. TUDCA-treated CKD-hMSCs, $$*p* < 0.01 vs. TUDCA-treated CKD-hMSCs pretreated with si-PRNP.

### TUDCA treatment of CKD-hMSCs increased renal proximal tubule epithelial cell (TH-1) viability in the CKD condition through upregulation of PrP^C^

To confirm the protective effect on TUDCA-treated CKD-hMSCs on the CKD condition induced by *P*-cresol, after TH-1 cells were exposed to *P*-cresol for 24 h during co-culture, we carried out co-culture of TH-1 cells with the upper chamber containing one of the following groups: Healthy-hMSCs, CKD-hMSCs, TUDCA-treated CKD-hMSCs (TUDCA+CKD- hMSC), TUDCA+CKD-hMSCs pretreated with si-*PRNP* (si-*PRNP*+CKD-hMSC), and TUDCA+CKD-hMSCs pretreated with si-*Scr* (si-*Scr*+CKD-hMSC) for 48 h. TH-1 cells showed reduced viability and decreased exrpression of PrP^C^ during exposure to *P*-cresol (Fig 5A, and B). Co-culture of TH-1 with the TUDCA+CKD-hMSC group increased cell viability to a level similar to that of the Healthy-hMSC group (Fig 5A). The TUDCA+CKD-hMSC group also showed increased expression of PrP^C^ in TH-1 cells (Fig 5B). These results showed that TH-1 viability may be increased by PrP^C^ and by the protective effect of TUDCA+CKD- hMSCs. To confirm that the TUDCA+CKD-hMSC group had greater mitochondrial membrane potential, MitoSOX analysis and ELISA were carried out and revealed reduced mitochondrial O_2_^•−^ concentration and enhanced complex I & IV activities (Fig 5C–E). In contrast, co-culture of TH-1 cells with si-*PRNP*+CKD-hMSC decreased cell viability and mitochondrial membrane potential. These results pointed to the key mechanism of the protective effect of TUDCA against CKD: an increase in mitochondrial membrane potential through upregulation of PrP^C^. These data indicated that TUDCA+CKD-hMSCs protect TH-1 cells against the CKD condition by increasing PrP^C^ expression and suggest that PrP^C^ plays a key role in the mechanism underlying TUDCA-enhanced mitochondrial membrane potential.

**Figure 5.**
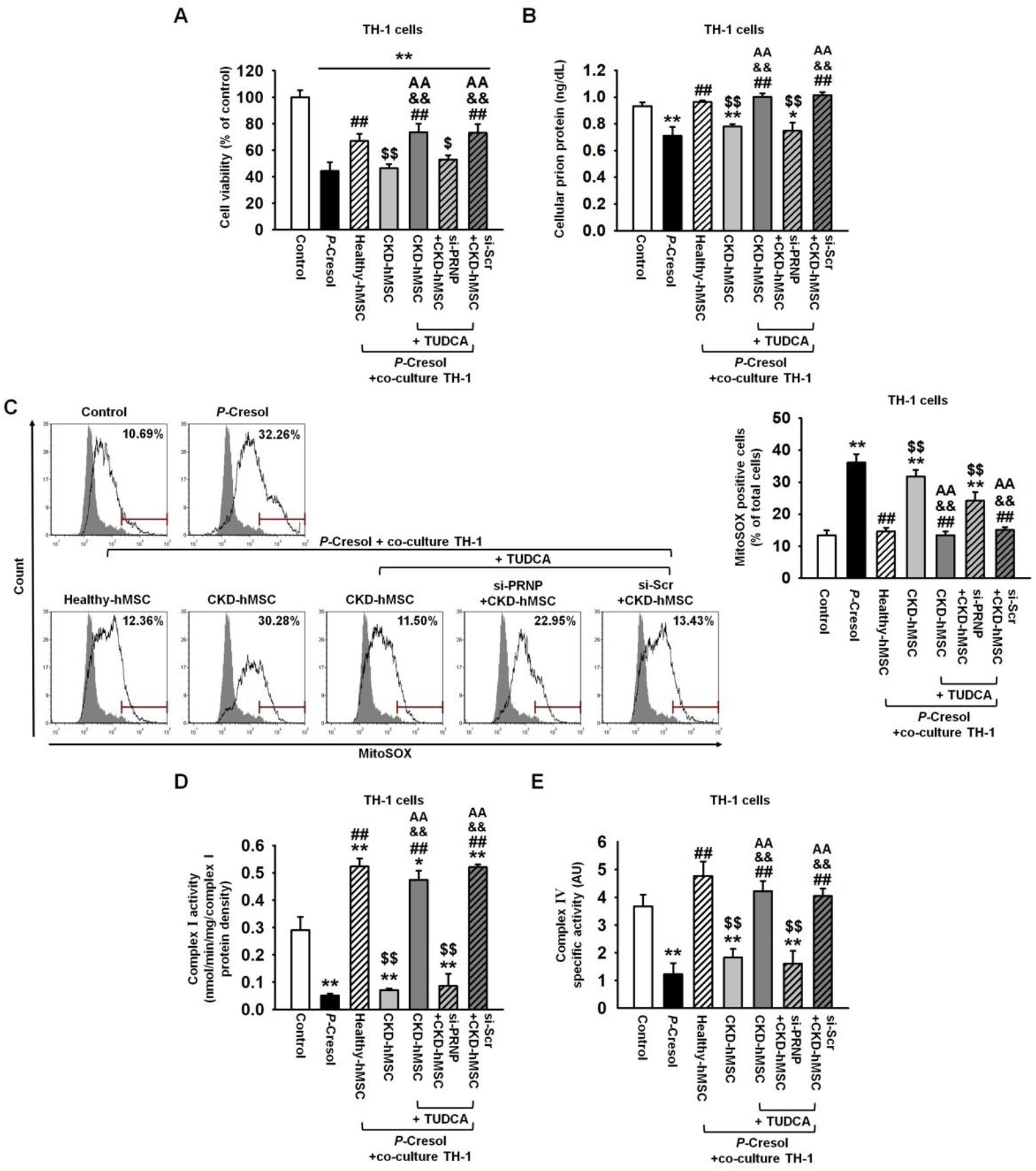
Human renal proximal tubular epithelial cells cocultured with TUDCA- treated CKD-hMSCs experience enhanced cell viability during *P*-cresol exposure. **A.** Treatment TH-1 cells with or without *P*-cresol cultured alone for 24 h, or cocultured with Healthy-hMSCs, CKD-hMSCs, TUDCA-treated CKD-hMSCs (TUDCA+CKD-hMSC), TUDCA-treated CKD-hMSCs pretreated with si-PRNP (si-PRNP+CKD-hMSC), or TUDCA-treated CKD-hMSCs pretreated with si-Scr (si-Scr+CKD-hMSC) using culture inserts for 48 h, after analysis of the proliferation of TH-1 cells. Values represent the mean ± SEM. \*\**p* < 0.01 vs. control, ##*p* < 0.01 vs. *P*-cresol group, $*p* < 0.05, and $$*p* < 0.01 vs. Healthy-hMSC group, && *p* < 0.01 vs. CKD-hMSC group, AA *p* < 0.01 vs. si-PRNP+CKD- hMSC group. **B.** Treatment of TH-1 cells with or without *P*-cresol cultured alone or with Healthy-hMSC group, CKD-hMSC group, TUDCA+CKD-hMSC group, si-PRNP+CKD-hMSC group, or si- Scr+CKD-hMSC group using culture inserts; we measured PrP^C^ expression only in lysates of TH-1 by ELISA. Values represent the mean ± SEM. \**p* < 0.05, and \*\**p* < 0.01 vs. control, ##*p* < 0.01 vs. *P*-cresol group, $$*p* < 0.01 vs. Healthy-hMSC group, && *p* < 0.01 vs. CKD- hMSC group, AA *p* < 0.01 vs. si-PRNP+CKD-hMSC group. **C.** The positive signals of MitoSOX were counted by FACS analysis with staining after treatment of TH-1 cells with or without *P*-cresol cultured alone, or with Healthy-hMSC group, CKD-hMSC group, TUDCA+CKD-hMSC group, si-PRNP+CKD-hMSC group, or si- Scr+CKD-hMSC group. Values represent the mean ± SEM. \*\**p* < 0.01 vs. control, ##*p* < 0.01 vs. *P*-cresol group, $$*p* < 0.01 vs. Healthy-hMSC group, && *p* < 0.01 vs. CKD-hMSC group, AA *p* < 0.01 vs. si-PRNP+CKD-hMSC group. **D, E**. Complex ? and complex ? activities were analyzed after treatment of TH-1 cells with or without *P*-cresol cultured alone, or with Healthy-hMSC group, CKD-hMSC group, TUDCA+CKD-hMSC group, si-PRNP+CKD-hMSC group, or si-Scr+CKD-hMSC group. Values represent the mean ± SEM. \**p* < 0.05, and\*\**p* < 0.01 vs. control, ##*p* < 0.01 vs. *P*- cresol group, $$*p* < 0.01 vs. Healthy-hMSC group, && *p* < 0.01 vs. CKD-hMSC group, AA *p* < 0.01 vs. si-PRNP+CKD-hMSC group.

### Increased serum PrP^C^ levels by TUDCA-treated CKD-hMSCs enhance recovery from kidney fibrosis through downregulation of inflammatory cytokines

We confirmed the CKD model in mice by feeding them with 0.25% adenine for 1 week, and then injected each group, Healthy-hMSCs, CKD-hMSCs, TUDCA-treated CKD-hMSCs (TUDCA+CKD-hMSC), TUDCA+CKD-hMSCs pretreated with si-*PRNP* (si-*PRNP*+CKD- hMSC), and si-*Scr*-pretreated TUDCA+CKD-hMSCs (si-*Scr*+CKD-hMSC) by intravenous (IV) tail injection on days 0, 5, and 10. At 25 days after discontinuation of adenine feeding, group TUDCA+CKD-hMSC had similar levels of BUN and creatinine compared with donor Healthy-hMSCs (Fig 6B and C). TUDCA+CKD-hMSCs also increased serum PrP^C^, which indicated increased repair of CKD damage to kidney cells through enhanced mitochondrial membrane potential (Fig 6D). Confirming aggravation of kidney fibrosis, a decrease of fibrosis on glomeruli and an increase of tubulo-interstitium were observed in mouse group TUDCA+CKD-hMSC according to H&E staining (Fig 6E). Kidney fibrosis could be aggravated by deficient IL-10, and by upregulation of TNF-α, a macrophage-secreted inflammation factor.(Sziksz et al, 2015) We confirmed by ELISA that injection of TUDCA+CKD-hMSCs into mice upregulated IL-10 and downregulation TNF-α in the serum of the mouse model of CKD (Fig 6F and G). In contrast, injection of si-*PRNP*+CKD-hMSC showed the opposite results, increasing BUN and creatinine levels (Fig 6B and C). These results indicated that the protective effect of TUDCA was attenuated by decreased serum PrP^C^ concentration (Fig 6D). Injection of si-*PRNP*+CKD-hMSC, as expected, downregulated IL-10 and upregulated TNF-α, which curtailed the increased kidney fibrosis and necrosis of kidney tissue (Fig 6E–G). There was a notable consequence: an increase in serum PrP^C^ by treatment with TUDCA enhanced CKD-hMSC stem cell therapy in the mouse model of CKD.

**Figure 6.**
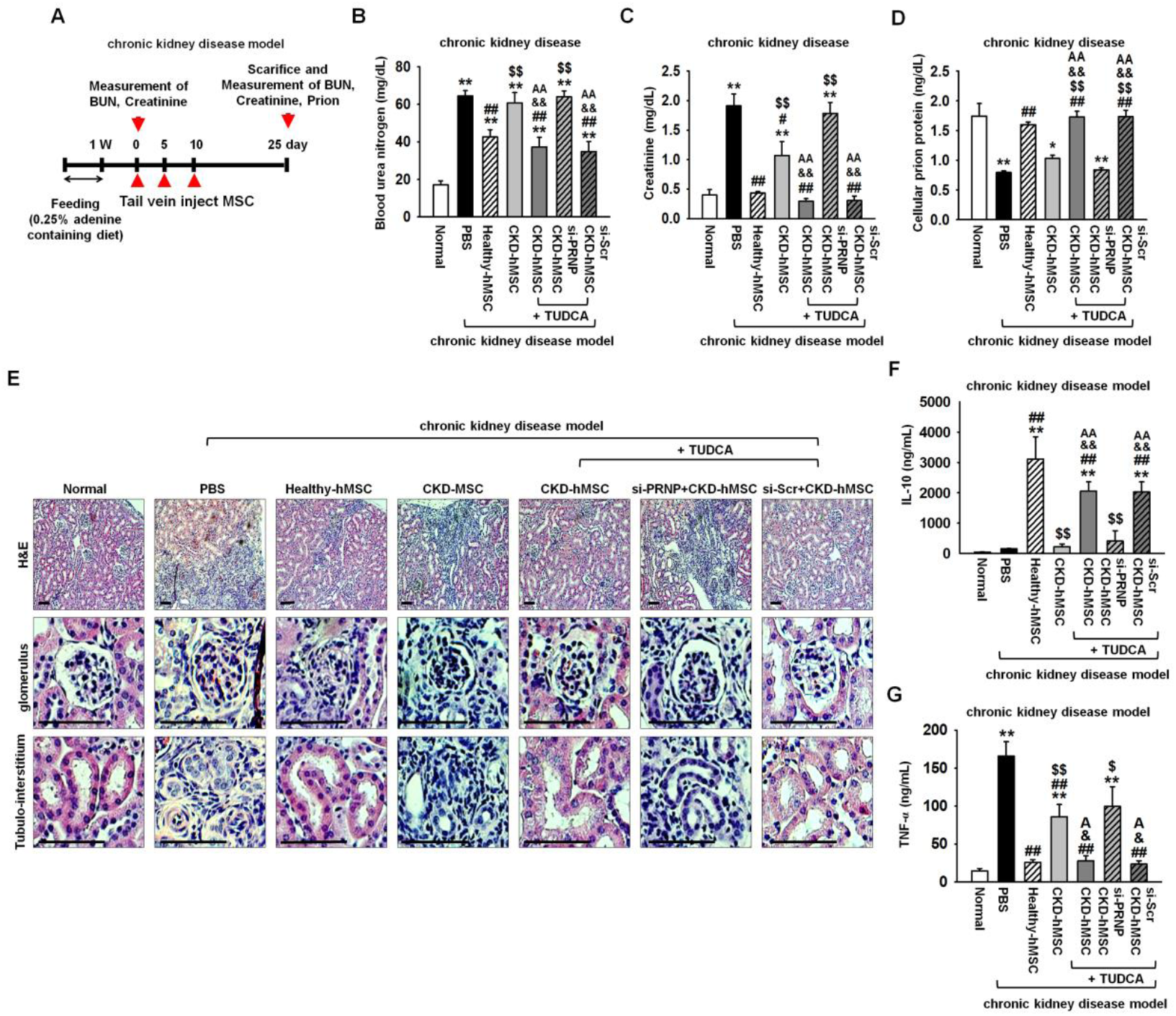
Transplantation of TUDCA-treated CKD-hMSCs protects against kidney fibrosis via upregulation of PrP^C^ in the mouse model of adenine-induced CKD. **A.** The scheme of the mouse model of CKD. The CKD model mice were fed 0.25% adenine for 1 week, and then we euthanized some mice in each group for validating the CKD model by measurement of BUN and creatinine. Each CKD mouse was injected with PBS or with Healthy-hMSCs, CKD-hMSCs, TUDCA-treated CKD-hMSCs (TUDCA+CKD-hMSC), TUDCA-treated CKD-hMSCs pretreated with si-PRNP (si-PRNP+CKD-hMSC), or TUDCA-treated CKD-hMSCs pretreated with si-Scr (si-Scr+CKD-hMSC) at 0, 5, and 10 days via a tail vein. After 25 days, measurement of BUN, creatinine, or PrP^C^ was carried out. **B, C, D.** At 25 days after discontinued feeding of adenine, we measured BUN, creatinine, and PrP^C^ in each mouse group in their serum samples (100 μL). Values represent the mean ± SEM. \**p* < 0.05, and \*\**p* < 0.01 vs. Normal, #*p* < 0.05, and ##*p* < 0.01 vs. PBS, $$*p* < 0.01 vs. Healthy-hMSC group, && *p* < 0.01 vs. CKD-hMSC group, AA *p* < 0.01 vs. si- PRNP+CKD-hMSC group. **B.** Regeneration of kidney tissue in CKD was confirmed by H&E staining. Scale bar = 100 μm. **F**, **G**. The level of IL-10 or TNF-α in each serum sample was analyzed by ELISA. Values represent the mean ± SEM. \*\**p* < 0.01 vs. Normal, ##*p* < 0.01 vs. PBS, $*p* < 0.05, and $$*p* < 0.01 vs. Healthy-hMSC group, & *p* < 0.05, and && *p* < 0.01 vs. CKD-hMSC group, A *p* < 0.05, and AA *p* < 0.01 vs. si-PRNP+CKD-hMSC group.

### Increased serum PrP^C^ concentration by TUDCA-treated CKD-hMSCs enhanced recovery from CKD-associated VD

After feeding of 0.25% adenine to mice for 1 week, we set up CKD-associated VD in mice by hindlimb ischemia surgery (Fig 7A). At postoperative day 3, injection with TUDCA+CKD-hMSC intravenously decreased levels of apoptosis-associated proteins— BAX, cleaved caspase 3, and cleaved PARP-1—and increased expression of neovascularization cytokines: hVEGF, hHGF, and hFGF (Fig 7B and C). These results indicated that TUDCA effectively treated CKD-associated VD through decreasing tissue apoptosis and upregulating neovascularization cytokines. At postoperative day 25, the blood perfusion ratio was analyzed by laser Doppler perfusion imaging and was found to be increased in the TUDCA+CKD-hMSC group to levels similar to those in the Healthy-hMSC group (Fig 7D and E). Moreover, the TUDCA+CKD-hMSC group showed a reduction in limb loss and foot necrosis (Fig 7F and G). To investigate neovascularization, we performed immunohistochemical staining for CD31 or α-SMA on postoperative day 25. Capillary density and arteriole density increased in the TUDCA+CKD-hMSC group (Fig 7H and I). By contrast, group si-*PRNP*+CKD-hMSC showed reduced neovascularization, according to measurement of neovascularization cytokines on postoperative day 3, and capillaries and arterioles had lower levels of CD31 and/or α-SMA on day 25, and increased necrosis of the foot, after we detected increased expression of apoptosis-associated proteins and decreased blood perfusion. These data indicated that TUDCA-treated CKD-hMSCs promoted neovascularization and functional recovery in kidney fibrosis-and-ischemia–injured tissue, and that regulation of mitochondrial PrP^C^ levels by TUDCA is important for the functionality of transplanted CKD-hMSCs in this tissue. These data indicated that TUDCA-treated CKD- hMSC enhanced neovascularization (through increased levels of PrP^C^) plays a pivotal role in the mouse model of CKD-associated VD.

**Figure 7.**
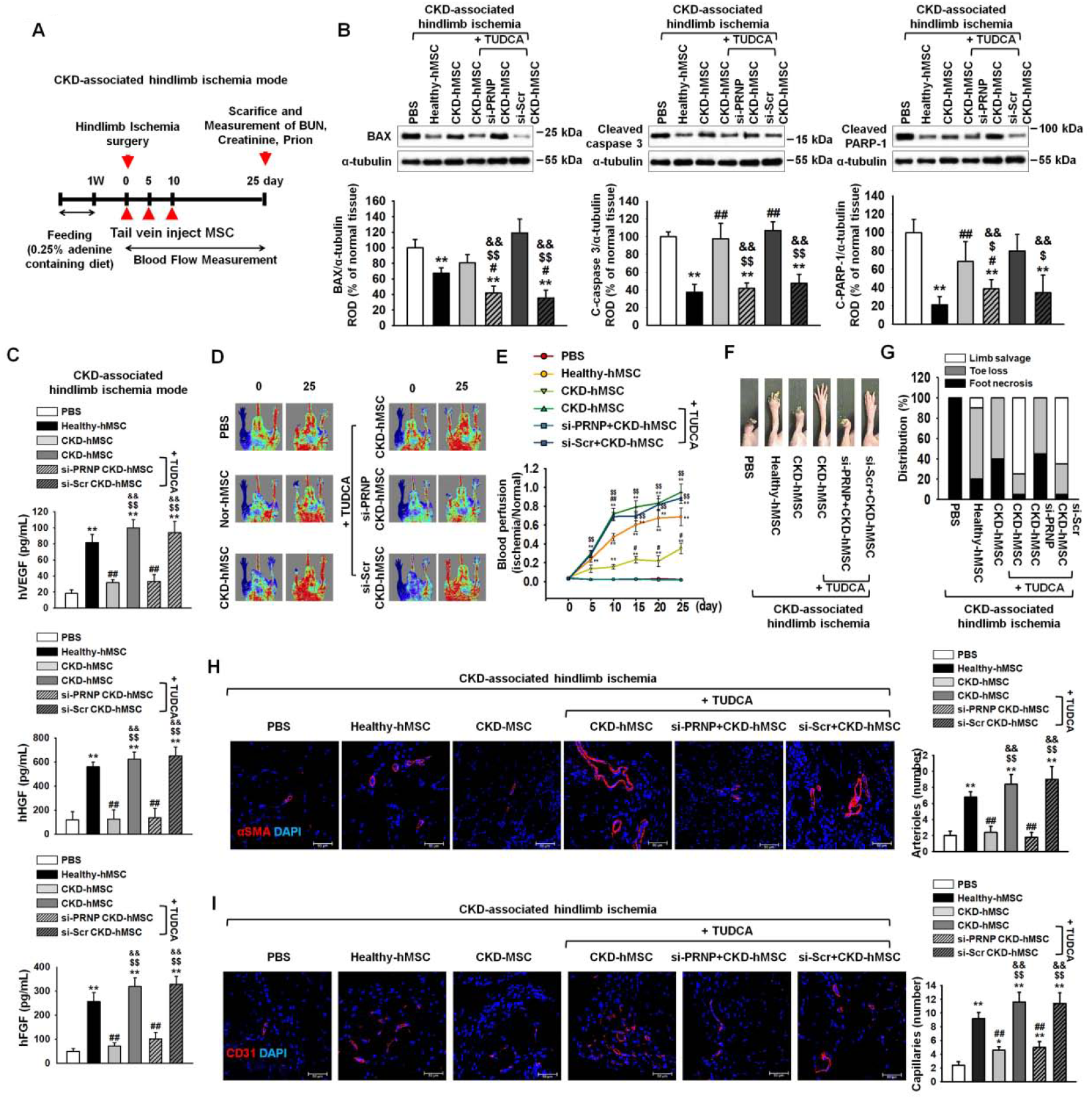
TUDCA-treated CKD-hMSCs enhanced neovascularization in the mouse model of CKD-associated hindlimb ischemia through upregulation of mitochondrial PrP^C^. **A.** The scheme of the mouse model of CKD-associated hindlimb ischemia. CKD-associated hindlimb ischemia mouse model was fed with 0.25% adenine for 1 week, and then we administered hindlimb ischemia in each mouse group surgically. Each CKD-associated hindlimb ischemia mouse model was injected with PBS or Healthy-hMSCs, CKD-hMSCs, TUDCA-treated CKD-hMSCs (TUDCA+CKD-hMSC), TUDCA-treated CKD-hMSCs pretreated with si-PRNP (si-PRNP+CKD-hMSC), or TUDCA-treated CKD-hMSCs pretreated with si-Scr (si-Scr+CKD-hMSC) at 0, 5, and 10 days via a tail vein. To measure tissue apoptosis and neovascularization at postoperative day 3, we euthanized some mice in each group. **B.** At postoperative day 3, western blot analysis quantified the cell apoptosis associated protein, BAX, cleaved caspase 3, and cleaved PARP-1 in each their hindlimb ischemia tissue group. The expression levels were determined by densitometry relative of α-tubulin. Values represent the mean ± SEM. \*\**p* < 0.01 vs. PBS, #*p* < 0.05 and ##*p* < 0.01 vs. Healthy-hMSC group, $*p* < 0.05, and $$*p* < 0.01 vs. CKD-hMSC, && *p* < 0.01 vs. si-PRNP+CKD-hMSC group. **C.** Expression of hVEGF, hHGF, and hFGF in each their hindlimb ischemia tissue lysis group was analyzed by ELISA. Values represent the mean ± SEM. \*\**p* < 0.01 vs. PBS, ##*p* < 0.01 vs. Healthy-hMSC group, $$*p* < 0.01 vs. CKD-hMSC, && *p* < 0.01 vs. si-PRNP+CKD- hMSC group. **D.** The CKD associated hindlimb ischemia mouse model improved in blood perfusion by laser Doppler perfusion imaging analysis of the ischemic-injured tissues of PBS, Healthy- hMSC, CKD-hMSC, TUDCA+CKD-hMSC, si-PRNP+CKD-hMSC, and si-Scr+CKD-hMSC at 0 day, and 25 days postoperation. **E.** The ratio of blood perfusion (blood flow in the left ischemic limb/blood flow in the right non-ischemic limb) was measured in each mouse of the five groups. Values represent the mean ± SEM. \*\**p* < 0.01 vs. PBS, #*p* < 0.05 and ##*p* < 0.01 vs. Healthy-hMSC group, $$*p* < 0.01 vs. CKD-hMSC. **F.** Representative image illustrating the different outcomes (foot necrosis, toe loss, and limb salvage) of CKD-associated ischemic limbs injected with five treatments at postoperative 25 days. **G.** Distribution of the outcomes in each group at postoperative day 25. **H**, **I.** At postoperative 25 days, capillary density, and arteriole density were analyzed by immunofluorescence staining for αSMA (Red) and CD31 (Red), respectively. Scale bar = 50 μm, 100 μm. Capillary density was quantified as the number of human αSMA- or CD31 positive cells, Values represent the mean ± SEM. \**p* < 0.05 and \*\**p* < 0.01 vs. PBS, ##*p* < 0.01 vs. Healthy-hMSC group, $$*p* < 0.01 vs. CKD-hMSC, && *p* < 0.01 vs. si-PRNP+CKD- hMSC group.

**Figure 8.**
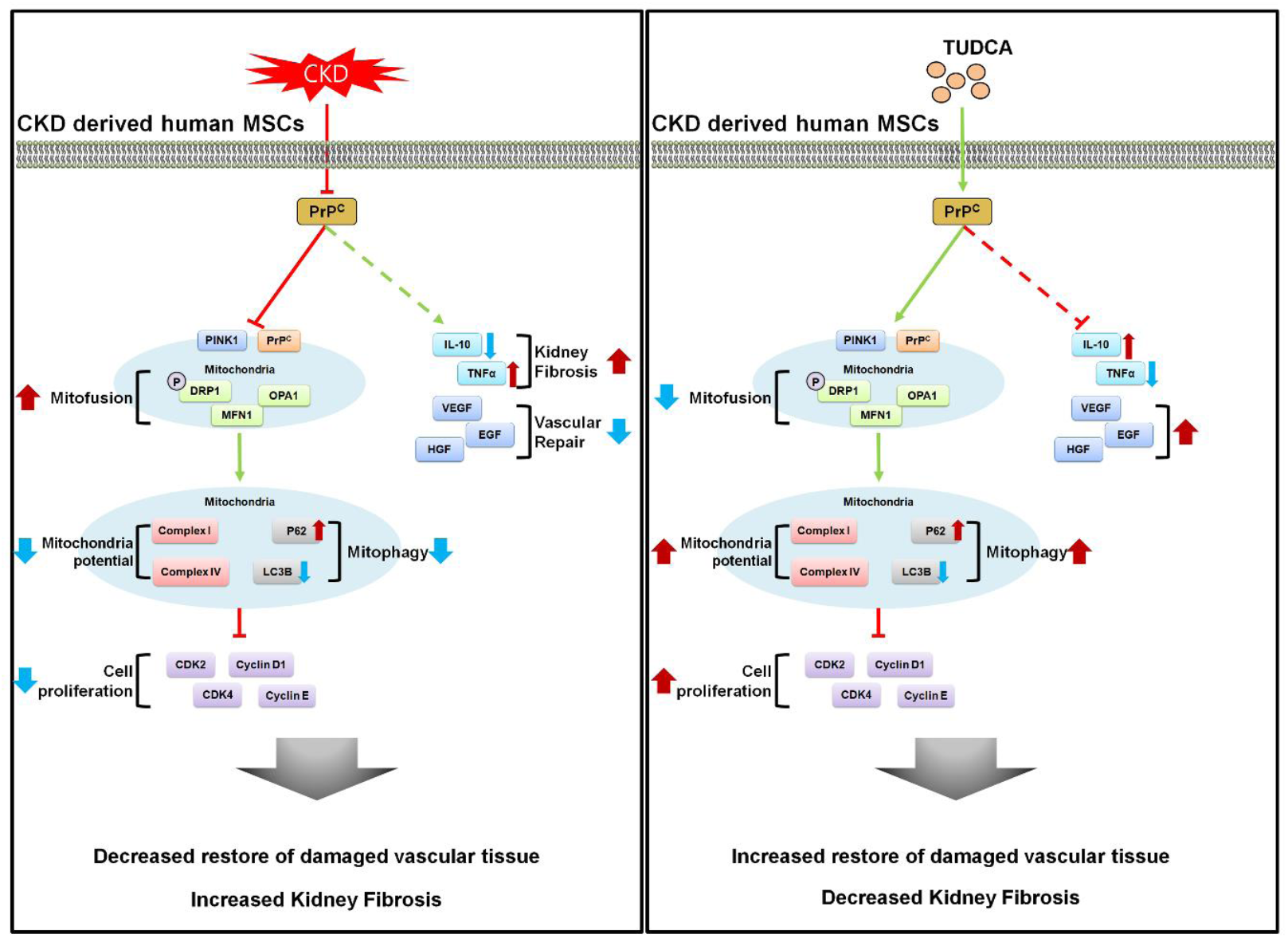
The scheme of autologous therapy with TUDCA-treated CKD-hMSCs: hMSC enhancement via increased concentration of PrP^C^. Reduction of mitochondrial functionality leads to decreased cellular proliferation and function, including decreased repair of damaged vascular tissue, ultimately unable to restore of kidney fibrosis. Increased in mitochondrial functionality leads to increased cellular proliferation, which enhances vascular repair of damaged cells. Healthier cellular state leads to increased autologous therapy of kidney fibrosis.

## Discussion

In our study, we reported that transplantation of TUDCA-treated patient-derived autologous MSCs reverses kidney fibrosis and increases neovascularization in a CKD VD model. In addition, we demonstrate that cellular PrP^C^ is a key molecule involved: we showed that a decreased level of PrP^C^ in the serum of the patients with CKD VD leads to reduced functionality of autologous MSCs derived from the patients with CKD. We have shown that TUDCA rescues MSCs derived from patients with CKD by increasing the amount of cellular prion protein (PrP^C^) in CKD-hMSCs, thus increasing the binding of PrP^C^ to PINK1 within mitochondria, restoring mitochondrial membrane potential in MSCs and inducing complex I & IV activation. In addition, we showed *in vivo* that the transplantation of TUDCA-treated CKD-hMSCs enhances therapeutic effects of MSC transplantation targeting CKD and associated conditions such as VD by increasing the PrP^C^ level. Our results suggest that therapeutic transplantation of CKD-hMSCs treated with TUDCA could lead to further studies as a potent therapy for patients who have both CKD and VD.

Cardiovascular disease (CD) such as peripheral ischemic disease is a common problem among patients with CKD, caused by hyperhomocysteinemia, oxidant stress, dyslipidemia, and elevated inflammatory markers after initiation of CKD.(Sarnak et al, 2003a; Sarnak et al, 2003b) We were able to determine that a CKD biomarker, eGFR (estimated glomerular filtration rate), is lower in patients’ serum at stages 3b–5 (average eGFR: 22.4), indicating that levels of BUN and creatinine circulate in blood serum. In addition, we discovered that levels of PrP^C^ are decreased in the serum of the patients with CKD-associated VD. One study has shown have shown that CKD in patients is accompanied by several mitochondrial aberrations such as high levels of ROS, inflammation, and a cytokine imbalance.(Schlieper et al, 2016) These results suggest that CKD-hMSCs possess decreased amounts of mitochondrial PrP^C^ (which binds to PINK1); this aberration causes mitochondrial dysfunction and damages cellular metabolism of CKD-hMSCs, potentially deprecating their therapeutic utility.

PrP^C^ is known to perform an integral function in bone marrow reconstitution, tissue regeneration, self-renewal, and recapitulation in stem cells.(Tanaka & Reddien, 2011; Zhang et al, 2006) PrP^C^ has been known to induce an abundant environment for stem cells to thrive, by providing nichelike conditions for stem cells to get activated and to proliferate.(Haddon et al, 2009; Martin-Lanneree et al, 2017; Zhang et al, 2006) In addition, our team has reported that the increased level of PrP^C^ inhibits oxidative stress, reducing cellular apoptosis through enhancement of SOD and catalase activity thereby increasing the therapeutic effectiveness in a hindlimb ischemia model.(Yoon et al, 2016) Similarly, our results suggested that TUDCA- treated CKD-hMSCs produce more PrP^C^, and that PrP^C^ binds to mitochondrial protein PINK1 as an integral pathway in TUDCA-initiated protection of CKD-hMSCs. Previous study proved PINK1 binds with TOM complex, thereby regulates of Parkin affects the mitochondria potential, and another study concluded mitochondrial PrP^C^ may be transported to TOM70.(Faris et al, 2017; Lazarou et al, 2012) Thus, we thought binging PrP^C^ with PINK1 might increase mitochondrial potential.

Another study revealed that downregulation of PINK1 disrupts the regulation of mitochondrial mitophagy in CKD, and mitochondria in the CKD model undergo impairment of mitochondrial fission thereby enlarged, dysfunctional mitochondria are formed.(Bueno et al, 2015; Forbes & Thorburn, 2018; Li et al, 2017) Impairment of mitochondrial fission causes cellular damage by decreasing the ability of mitochondria to remove damaged components; this observation suggests that PINK1-mediated mitochondrial fission is critical for healthy renal function.(Forbes & Thorburn, 2018; Zhan et al, 2013) In our study, we demonstrated that the TUDCA-treated CKD-hMSCs have greater mitofission according to western blot analysis of mitofission-inhibitory protein p-DPR1 (phosphorylation of Ser637) and mitofusion proteins MFN1 and OPA1 as well as increased stabilization of mitochondrial membrane potential as well as a decrease in the number of abnormal mitochondria. More specifically, we found that PrP^C^ binding to PINK1 is crucial for retaining stability of the inner mitochondrial membrane and for enhancing the mitochondrial membrane potential according to TEM imaging. These results suggest that the mitochondrial dysfunction in CKD-hMSCs is exacerbated due to decreased stability of PINK1 bound to PrP^C^, and that TUDCA counters these limitations by protecting CKD-hMSCs via upregulation of PrP^C^, thereby repairing downstream mitochondrial membrane potential and cellular proliferation. Our results of the BrdU, cell cycle, and CDK4/Cyclin D1 assays indicate that TUDCA+CKD-hMSCs could potentially rescue a human damaged kidney via enhanced PrP^C^ levels and increased mitochondrial functionality.

We aimed to verify in CKD-hMSCs that TUDCA enhanced mitochondrial membrane potential and increased the amount of PrP^C^, which has protective effects, thus possibly recovering other damaged kidney cells through TUDCA-treated CKD-hMSCs using human renal proximal tubular epithelial cells (TH-1). We developed a CKD conditional model where TH-1 cells are exposed to *P*-cresol for 24 h, and then seeded in the upper chamber for co-culture with respective MSC groups for 48 h. The TUDCA+CKD-hMSCs group, as expected, increased TH-1 cell viability by increasing PrP^C^. Therefore, group TUDCA+CKD-hMSCs enhanced mitochondrial membrane potential, reduced mitochondrial O_2_^•−^, and increased complex I & IV activities, similar to the protective effect of TUDCA. These data support our theory, that effective upregulation of PrP^C^ could lead to regulate mitochondrial membrane potential and that despite the use of damaged cells, stem cell therapy can be enhanced by treatment with TUDCA. In contrast, co-culture of TH-1 cells with si-*PRNP*+CKD-hMSCs group decreased viability of TH-1 cells, because the impossible uptake of PrP^C^ blocked the protective effect of TUDCA. Therefore, group si-*PRNP*+CKD-hMSCs increased mitochondrial O_2_^•−^ levels and decreased complex I & IV activity. We verified that TUDCA not only increased cell proliferation through enhancement of mitochondrial membrane potential but also restored CKD-damaged kidney tubular cells via an increased level of PrP^C^.

Finally, we showed *in vivo* that TUDCA-treated CKD-hMSCs are a potent tool for tackling CKD and for combating its associated kidney fibrosis and neovascularization in a mouse model of CKD-associated hindlimb ischemia. This is because TUDCA-treated CKD- hMSCs overcome the pathophysiological conditions associated with the host interaction. We confirmed CKD mice after feeding 0.25% adenine for 1 week, and then set up a hindlimb ischemic mouse model at 1 week after discontinuing adenine feeding for stability of the mouse model. Each hMSC group injected intravenously via tail injection on days 0, 5, and 10. First, at 25 days after discontinuation of adenine feeding, we confirmed in the serum of the TUDCA+CKD-hMSCs group that the levels of kidney dysfunction markers—BUN and creatinine—and PrP^C^ in serum, were normalized. We directly examined the restored kidneys of the mice through H&E staining that CKD fibrosis and necrosis decreased in teh TUDCA+CKD-hMSC–injected group. H&E staining showed that injected TUDCA+CKD- hMSCs decreased fibrosis, reduced glomerulus swelling, and increased tubule interstitium. Other studies have suggested that increased inflammatory factors are crucial for determining the development of renal dysfunction and fibrosis.(Rodell et al, 2015; Sziksz et al, 2015) In our mouse model of CKD-associated hindlimb ischemia, injected TUDCA+CKD-hMSCs caused significant changes in the inflammatory cytokines such as increased levels of IL-10 and decreased levels of TNF-α, which suggest that inflammation was lowered. In addition, we aimed to evaluate similar therapeutic effects at the ischemic injury site of the mice. We observed that injected TUDCA+CKD-hMSCs in a mouse group saved vascular functions via downregulation of apoptosis-related proteins and upregulation of neovascularization proteins on postoperative day 3, with the increased blood perfusion ratio, limb salvage, and vessel formation, such as development of arteries and capillary vessels on postoperative day 25 in a PrP^C^-dependent pathway. Injection of si-*PRNP*+CKD-hMSCs increased kidney fibrosis and thereby induced a decrease of IL-10 and increase of TNF-α levels and attenuated the functional recovery and neovascularization in the mouse model of CKD-associated hindlimb ischemia by blocking expression of PrP^C^.

In our study, we demonstrated a novel finding that CKD patient–derived MSCs manifest reduced functionality because of the decreased level of PrP^C^ in serum. In addition, we propose that TUDCA effectively potentiates autologous CKD-hMSCs to exert therapeutic effects by targeting CKD-associated hindlimb ischemia in the mouse model and a fibrotic cell population, thus overcoming the obstacles of compromised stem cell function of CKD- hMSCs. We also demonstrated the mechanistic pathway via which TUDCA enhanced mitochondrial membrane potential: by enhancing the expression of PrP^C^ thereby promoting the binding of PrP^C^ to PINK1 in the stem cells and increasing the supply of blood serum PrP^C^. These results suggest that TUDCA-treated CKD-hMSCs could offer a novel approach to targeting CKD-associated hindlimb ischemia via the use of a safe, feasible autologous cell source via intravenous injection of protected MSCs into the patients via precise regulation of PrP^C^.

## Materials and Methods

### Serum of patients with CKD and the healthy control

The study was approved by the local ethics committee, and informed consent was obtained from all the study subjects. Explanted serum (n = 25; CKD-patient) and the healthy control fulfilling transplantation criteria (n = 10; Healthy control) were obtained from the Seoul national hospital in Seoul, Korea. (IRB: SCHUH 2018-04-035). All CKD diagnoses wasd defined as abnormalities of kidney function as measure estimated glomerular filtration rate (eGFR) < 25 ml/min/1.73m^2^ over 3 months (stage 3b∼5).

### Human Healthy MSCs and human CKD MSCs cultures

Human adipose tissue-derived Healthy-hMSCs (n = 4) and CKD-hMSCs (n = 4) were obtained from Soonchunhyang University Seoul hospital (IRB: SCHUH 2015-11-017). We defined CKD as impaired kidney function according to estimated glomerular filteration rate [eGFR] <35 ml/(min·1.73 m^2^) for more than 3 months (stage 3b). The supplier certified the expression of MSC surface positive markers CD44, and Sca-1, and negative marker CD45, and CD11b. When cultured in specific differentiation media, MSCs also differentiate into chondrogenic, adipogenic, and osteogenic cells. Healthy hMSCs and CKD-hMSCs were cultured in α-Minimum Essential Medium (α-MEM; Gibco BRL, Gaithersburg, MD, USA) supplemented with 10% (v/v) of fetal bovine serum (FBS; Gibco BRL) and a 100 U/ml penicillin/streptomycin (Gibco BRL). Healthy-hMSCs and CKD-hMSCs were grown in a humidified 5% CO_2_ incubator at 37 °C.

### Detection of PrP^C^ in serum or cell media

Concentrations of PrP^C^ in the human healthy control group or CKD patient group, or in a cell lysate of untreated Healthy-hMSCs or CKD-hMSCs with TUDCA, or in Healthy-hMSCs or CKD-hMSCs treated with TUDCA were determined with a commercially available ELISA kit (Lifespan Biosciences, Seattle, WA, USA). Next, 100 μl from each serum group or cell media group was used for these experiments. Triplicate measurements were performed in all ELISAs. Expression levels of PrP^C^ were quantified by measuring absorbance at 450 nm using a microplate reader (BMG Labtech, Ortenberg, Germany).

### Mitochondrial-fraction preparation

Harvested Healthy-hMSCs or CKD-hMSCs were lysed in mitochondria lysis buffer with vortexing and were subsequently incubated for 10 min on ice. The cell lysates were centrifugated at 3000 rpm at 4 °C for 5 min, and the supernatant was collected as a cytosol fraction. The residual pellet was lysed with RIPA lysis buffer and then centrifuged at 15,000 rpm and 4 °C for 30 min.

### Western blot analysis

The whole-cell lysates, cytosol fraction lysates, or mitochondrial lysates from Healthy- hMSCs or CKD-hMSCs (30 μg protein) were separated by sodium dodecyl sulfate-polyacrylamide gel electrophoresis (SDS-PAGE) in a 8–12% gel, and the proteins were transferred to a nitrocellulose membrane. After the blots were washed with TBST (10 mM Tris-HCl [pH 7.6], 150 mM NaCl, 0.05% Tween 20), the membranes were blocked with 5% skim milk for 1 h at room temperature and incubated with the appropriate primary antibodies: against PrP^C^, MFN1, or β-actin (Santa Cruz Biotechnology,llas, TX, USA), PINK1, OPA1, P62, LC3B, and VDAC1 (NOVUS, Littleton, CT, USA), and p-DPR1 (Cell signaling, Danvers, MA, USA). The membranes were then washed, and the primary antibodies were detected by means of goat anti-rabbit IgG or goat anti-mouse IgG antibodies (conjugated secondary antibodies) (Santa Cruz Biotechnology). The bands were detected by enhanced chemiluminescence (Amersham Pharmacia Biotech, Little Chalfont, UK).

### Immunoprecipitation

The mitochondrial fraction of Healthy-hMSCs or CKD-hMSCs was lysed with lysis buffer (1% Triton X-100 in 50 mM Tris-HCl [pH 7.4], containing 150 mM NaCl, 5 mM EDTA, 2 mM Na_3_VO_4_, 2.5 mM Na_4_PO_7_, 100 mM NaF, and protease inhibitors). Mitochondrial lysates (200 μg) were mixed with an anti-PINK1 antibody (Santa Cruz Biotechnology). The samples were mixed with the Protein A/G PLUS-Agarose Immunoprecipitation Reagent (Santa Cruz Biotechnology) at 4°C for incubation over 4 h, and incubated additionally at 4°C for 12 h. The beads were washed four times, and the bound protein was released from the beads by boiling in SDS-PAGE sample buffer for 7 min. The precipitated proteins were analyzed by western blotting with an anti-PrP^C^ antibody (Santa Cruz Biotechnology).

### Measurement of mitochondrial O_2_^•−^ production

To measure the formation of mitochondrial O_2_^•−^, the mitochondrial superoxide of Healthy- hMSCs or CKD-hMSCs was measured by means of MitoSOX™ (Thermo Fisher Scientific, Waltham, MA, USA). These cells were trypsinized for 5 min and then centrifuged at 1200 rpm for 3 min, washed with PBS twice and then incubated with a 10 μM MitoSOX™ solution in PBS at 37°C for 15 min. After that, the cells were washed more than 2 times with PBS. Next, we resuspended these cells in 500 μL of PBS, and then detected the signals with MitoSOX™ by fluorescence-activated cell sorting (FACS; Sysmex, Kobe, Japan). Cell forward scatter levels were determined in MitoSOX™-positive cells, analyzed by means of the Flowing Software (DeNovo Software, Los Angeles, CA, USA).

### Cell cycle analysis

Healthy-hMSCs, TUDCA-treated Healthy-hMSCs, CKD-hMSCs, TUDCA-treated CKD- hMSCs, TUDCA-pretreated CKD-hMSCs after treatment with si-PRNP, or TUDCA- pretreated CKD-hMSCs after treatment with si-Scr were harvested and fixed with 70% ethanol at −20 °C for 2 h. After two washes with cold PBS, the cells were subsequently incubated with RNase and the DNA-intercalating dye propidium iodide (PI; Sysmex) at 4 °C for 1 h. Cell cycle of the PI-stained cells was characterized by FACS (Sysmex). Events were recorded for at least 10^4^ cells per sample. The sample data were analyzed in the FCS express 5 software (DeNovo Software). Independent experiments were repeated 3 times.

### Electron microscopy

For this purpose, the cells were fixed in 3% glutaraldehyde and 2% paraformaldehyde in 0.1 M sodium cacodylate buffer at pH 7.3. Morphometric analyses (the number of mitochondria per cell and mitochondrial size) were performed in ImageJ software (NIH; version 1.43). At least 10 cells from low-magnification images (×10,000) were used to count the number of mitochondria per hMSCs (identified by the presence of lamellar bodies). At least 100–150 individual mitochondria, from 3 different lungs per group at high magnification (×25,000 and ×50,000), were used to assess the perimeter and the area.

### Cell proliferation assay

Cell proliferation was examined by a 5-bromo-2′-deoxyuridine (BrdU) incorporation assay. MSCs were cultured in 96-well culture plates (3,000 cells/well). MSCs were exposed to Cripto (0, 1, 10, 50, 100, or 200 ng/ml) for a fixed period of 24 h or to 100 ng/ml Cripto for 12, 24, or 48 h. BrdU incorporation into newly synthesized DNA of proliferating cells was assessed by an ELISA colorimetric kit (Roche, Basel, Swiss). To perform the ELISA, 100 μg/ml BrdU was added to MSC cultures and incubated at 37°C for 3 h. An anti-BrdU antibody (100 μL) was added to MSC cultures and incubated at room temperature for 90 min. After that, 100 μL of a substrate solution was added and 1 M H_2_SO_4_ was employed to stop the reaction. Light absorbance of the samples was measured on a microplate reader (BMG Labtech) at 450 nm.

### Kinase assays of complex I & IV activities and CKD4/Cyclin D1 activity

The cells were lysed with RIPA lysis buffer (Thermo Fisher Scientific). Activity of complex I & IV and CDK4 kinase assays were performed by means of each the CKD4 Kinase Assay Kit (Cusabio, Baltimore, USA) and complex I & IV assays (abcam, Cambridge, UK). 30-50 μg of total cell lysates was subjected to these experiments. Activation of complex I & IV and CDK4 kinase assays quantified by measuring absorbance at 450 nm on a microplate reader (BMG).

### Co-culture of human renal proximal tubular epithelial cells with Healthy-hMSCs or CKD-hMSCs

Healthy-hMSCs or CKD-hMSCs and THP-1 cells were co-cultured in Millicell Cell Culture Plates (Millipore, Billerica, MA, USA), in which culture media are in indirect contact with cells. TH-1 cells were seeded in the lower compartments, and then exposed to *P*-cresol for 24 h, and then Healthy-hMSCs, CKD-hMSCs, TUDCA-treated CKD-hMSCs (TUDCA+CKD- hMSC), TUDCA-pretreated CKD-hMSCs treated with si-PRNP (si-PRNP+CKD-hMSC), or TUDCA-pretreated CKD-hMSCs treated with si-Scr (si-Scr+CKD-hMSC) were seeded onto the Transwell membrane inserts for incubation for 48 h. They were incubated in a humidified atmosphere containing 5% of CO_2_ at 37 °C. TH-1 cell proliferation, expression of PrP^C^, MitoSOX, and complex I & IV activities were assessed as described above for the proliferation assay, ELISAs, and FACS analysis.

### Detection of PrP^C^ in cell lysates

Concentrations of PrP^C^ in TH-1 cells incubated with or without *P*-cresol alone or co-cultured with one of the following cell types—Healthy-hMSCs, CKD-hMSCs, TUDCA+CKD-hMSCs, si-PRNP+CKD-hMSCs, or si-Scr+CKD-hMSCs—were determined using a commercially available ELISA kit (Lifespan Biosciences). 30 μg of total protein from each group of TH-1 cell lysates was subjected to these experiments. Triplicate measurements were performed in all ELISAs. Expression levels of PrP^C^ were quantified by measuring absorbance at 450 nm on a microplate reader (BMG Labtech).

### Ethics statement

All animal care procedures and experiments were approved by the Institutional Animal Care and Use Committee of Soonchunhyang University Seoul Hospital (IACUC2013-5) and were performed in accordance with the National Research Council (NRC) Guidelines for the Care and Use of Laboratory Animals. The experiments were performed on 8-week-male BALB/c nude mice (Biogenomics, Seoul, Korea) maintained on a 12-h light/dark cycle at 25 °C in accordance with the regulations of Soonchunhyang University Seoul Hospital.

### The CKD model

Eight-week-old male BALB/c nude mice were fed an adenine-containing diet (0.25% adenine in diet) for 1 to 2 weeks.(Lin et al, 2017) Mouse body weight was measured every week. The mice were randomly assigned to 1 of 4 groups consisting of 10 mice in each group. After euthanasia, blood was stored at −80 °C for measurement of blood urea nitrogen (BUN) and creatinine.

### The murine hindlimb ischemia model

To induce vascular disease and to assess neovascularization in the mouse model of CKD, after adenine-loaded feeding for 1 week, the murine hind limb ischemia model was established as previously described(Limbourg et al, 2009) with minor modifications. Ischemia was induced by ligation and excision of the proximal femoral artery and boundary vessels of the CKD mice. No later than 6 h after the surgical procedure, PBS, CKD-hMSCs, TUDCA-treated CKD-hMSCs, TUDCA-pretreated CKD-hMSCs treated with si-PRNP, or TUDCA-pretreated CKD-hMSCs treated with si-Scr in PBS were intravenously injected into a tail vein (10^6^ cells per 100 μL of PBS per mouse; 5 mice per treatment group) of CKD mice. Blood perfusion was assessed by measuring the ratio of blood flow in the ischemic (left) limb to that in the nonischemic (right) limb on postoperative days 0, 5, 10, 15, 20, and 25 by laser Doppler perfusion imaging (LDPI; Moor Instruments,Wilmington, DE).

### Hematoxylin and eosin (H&E), and immunohistochemical staining

At 25 days after the operation, the ischemic thigh tissues were removed and fixed with 4% paraformaldehyde (Sigma), and each tissue sample was embedded in paraffin. For histological analysis, the samples were stained with H&E in kidney tissues to determine fibrosis and histopathological features, respectively. Immunofluorescent staining was performed with primary antibodies against CD31 (Santa Cruz Biotechnology), and α-SMA (alpha-smooth muscle actin; Santa Cruz Biotechnology), followed by secondary antibodies conjugated with Alexa Fluor 488 or 594 (Thermo Fisher Scientific). Nuclei were stained with 4′,6-diaminido-2-phenylindol (DAPI; Sigma), and the immunostained samples were examined by confocal microscopy (Olympus, Tokyo, Japan).

### Detection of human growth factors

Concentrations of VEGF, FGF, and HGF in hindlimb ischemia–associated CKD tissue lysates were determined with commercially available ELISA kits (R&D Systems, Minneapolis, MN, USA). On postoperative day 3, 300 μg of total protein from hindlimb ischemia tissue lysates was subjected to these experiments. Triplicate measurements were carried out in all the ELISAs. Expression levels of growth factors were quantified by measuring absorbance at 450 nm on the microplate reader (BMG Labtech).

### RNA isolation and reverse transcription polymerase chain reaction

To isolate RNA, Heathy-hMSCs and CKD-hMSCs were extracted in TRIzol^®^ reagent (Invitrogen, Carlsbad, CA, USA). The concentration of RNA, of which quality was assessed using the 260/280 nm absorbance ratio, was quantified using a microplate reader (Tecan Group AG, Mannedorf, Switzerland) and reverse transcription polymerase chain reaction (RT-PCR) was performed with 100 ng total RNA by RevertAid First Strand cDNA Synthesis Kit (Thermo Fisher Scientific, Waltham, MA, USA). PCR amplification was performed with synthetic gene-specific primer for Osteopontin (OPN) forward, 5’- CTCCATTGACTCGAACGACTC-3’; OPN reverse, 5’- CAGGTCTGCGAAACTTCTTAGAT-3’ FABP4 forward, 5’- CGTGGAAGTGACGCCTTTCATG-3’; FABP4 reverse, 5’- ACTGGGCCAGGAATTTGACGAA-3’; SOX9 forward, 5’- AGCGAACGCACATCAAGAC-3’; SOX9 reverse, 5’-CTGTAGGCGATCTGTTGGGG-3’; β-actin forward, 5’- AACCGCGAGAAGATGACC–3’; β-actin reverse, 5’- AGCAGCCGTGGCCATCTC-3’.

### Quantitative real-time PCR

Quantitative real-time PCR (qRT-PCR) analysis was analyzed Maxima SYBR Green/ROX qPCR Master Mix (Thermo Fisher Scientific). qRT-PCR reaction using the PIKOREAL 96 (Thermo Fisher Scientific) was performed under cycling conditions: 95 °C for 15 seconds (denaturation), 55 °C for 30 seconds (annealing), and 72 °C for 60 seconds (extension) for 45 cycles. The gene expression level, normalized to β-actin, was then calculated using the 2-ΔΔCt formula with reference to the hMSC. Primer sequences were as follows: OPN forward, 5’-CTCCATTGACTCGAACGACTC-3’; OPN reverse, 5’- CAGGTCTGCGAAACTTCTTAGAT-3’ FABP4 forward, 5’- CGTGGAAGTGACGCCTTTCATG-3’; FABP4 reverse, 5’- ACTGGGCCAGGAATTTGACGAA-3’; SOX9 forward, 5’- AGCGAACGCACATCAAGAC-3’; SOX9 reverse, 5’-CTGTAGGCGATCTGTTGGGG-3’; β-actin forward, 5’- AACCGCGAGAAGATGACC–3’; β-actin reverse, 5’- AGCAGCCGTGGCCATCTC-3’.

### MSC differentiation

To test the differentiation of Healthy-hMSCs and CKD-hMSCs, cells were grown in StemPro adipogenic, osteogenic, or chondrogenic culture medium (Thermo Fisher Scientific). Cells were grown in the adipogenic medium for 1 week, osteogenic medium for 1 weeks, and chondrogenic medium for 2 weeks. Adipocytes were stained with oil red O (Sigma-Aldrich, St. Louis, MO, USA) for 10 min, osteoblasts were stained with alkaline phosphatase stain kit (Sigma-Aldrich) for 10 min, and chondrocytes were stained with Safranin O (Sigma-Aldrich) for 5 min. The samples were observed by inverted microscopy (Nikon, Tokyo, Japan).

### SOD Activity

Heathy-hMSCs and CKD-hMSCs were harvested from the culture dish by scraping with a rubber policeman on ice. The cells were extracted using extraction buffer. Cell lysates (50 µg) were allowed to react with superoxide dismutase, immediately after which signals were measured each minute for 15 min by ELISA reader (BMG labtech, Ortenberg, Germany) at 450 nm.

### Catalase Activity

Heathy-hMSCs and CKD-hMSCs protein extracts (40 µg) were incubated with 20 mM H_2_O_2_ for 30 min. Next, 50 mM Amplex Red reagent (Thermo Fisher Scientific) and 0.2 U/mL of horseradish peroxidase (Sigma-Aldrich) were added and incubated for 15 min at RT. Changes in the absorbance values associated with H_2_O_2_ degradation were measured by ELISA reader (BMG Labtech) at 563 nm.

### Statistical analysis

Results were expressed as the mean ± standard error of the mean (SEM) and evaluated by Two-tailed Student’s t test was used to compute the significance between the groups, or one- or two- way analysis of variance (ANOVA). Comparisons of three or more groups were made by using Dunnett’s or Tukey’s post-hoc test. Data were considered significantly different at P < 0.05.

## Author contributions

Y.M.Y. contributed to acquisition of data, analysis and interpretation of data, and drafting of the manuscript. S.M.K, Y.S.H, C.W.Y, and J.H.L provided interpretation of data, statistical analysis, and drafting of the manuscript. H.J.N. developed the study concept and drafting of the manuscript. S.H.L. developed the study concept and design, and performed acquisition of data, analysis and interpretation of data, drafting of the manuscript, procurement of funding, and study supervision.

## Acknowledgements

This work was supported by the Soonchunhyang University Research Fund, a National Research Foundation grant funded by the Korean government (NRF- 2017M3A9B4032528). The biospecimens for this study were provided by the Seoul National University Hospital Human Biobank, a member of the Korea Biobank Network, which is supported by the Ministry of Health and Welfare. All samples derived from the National Biobank of Korea were obtained with informed consent under institutional review board-approved protocols. The funders had no role in the study design, data collection or analysis, the decision to publish, or preparation of the manuscript.

## Expanded View Figures legends

**Figure EV1.**
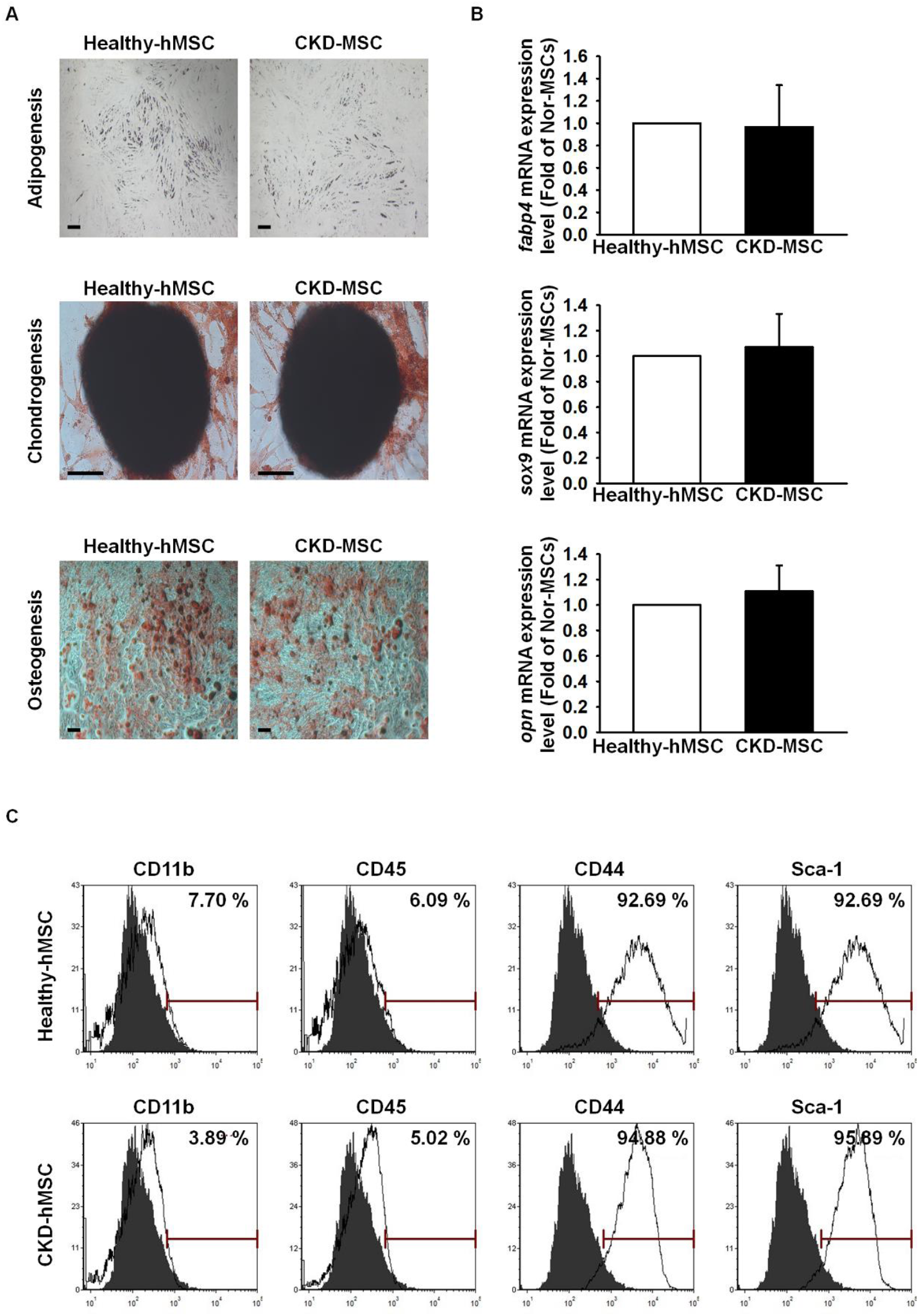
Healthy-hMSCs and CKD-hMSCs did not change cell differentiation. **A.** Healthy-hMSC and CKD-hMSC possible cell differentiation adipogenesis, chondrogenesis, and osteogenesis. Scale bar = 100 μm. **B.** Healthy-hMSCs and CKD-hMSCs retained similar expression level of differentiation association mRNA, *fabp4*, *sox9*, and *opn*. **C.** Healthy-hMSC and CKD-hMSC measured stem cell surface marker CD11b, and CD45, negative marker, and CD44, and Sca-1, positive marker.

**Figure EV2.**
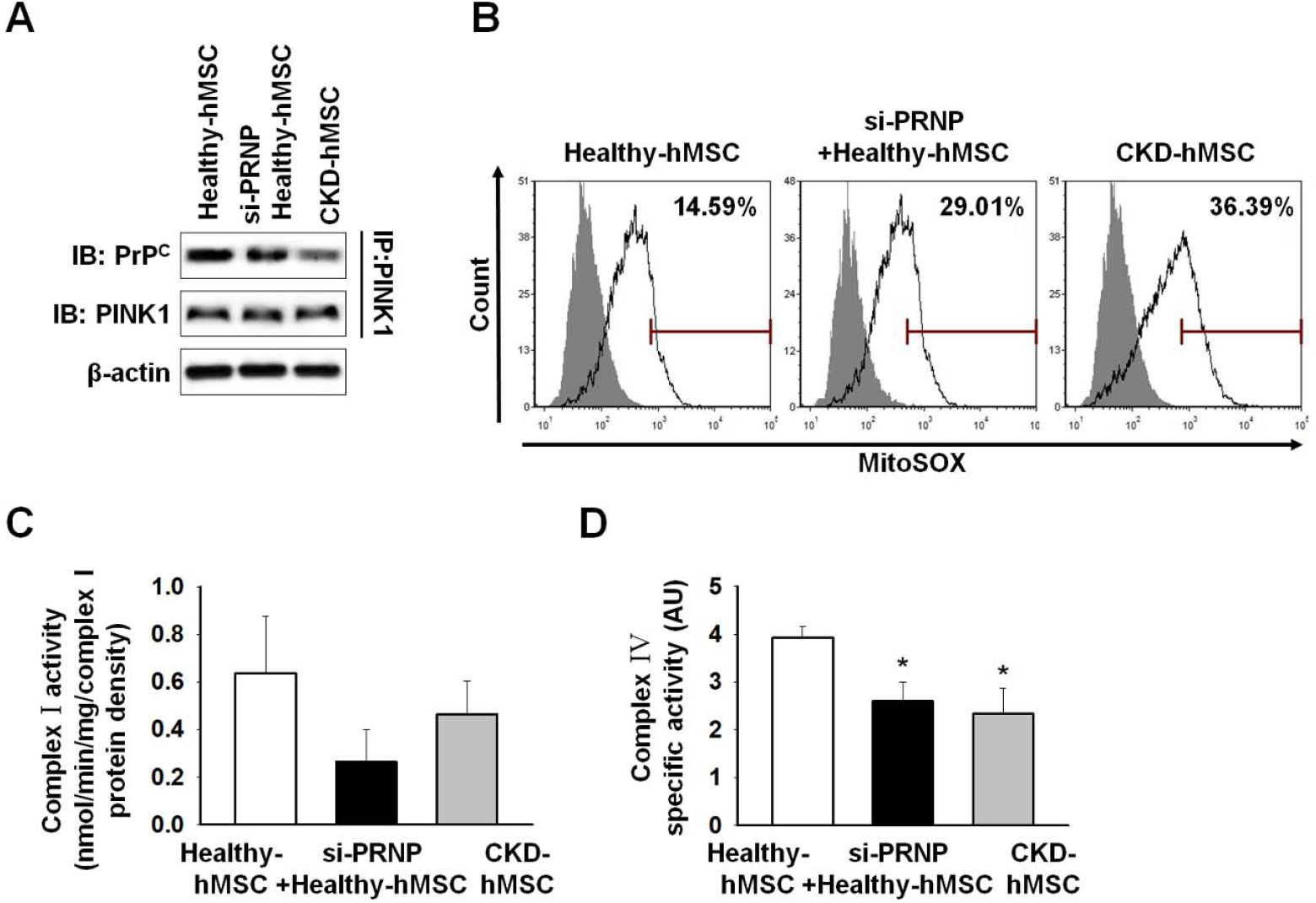
Treatment of Healthy-hMSCs with si-PRNP decreased mitochondrial membrane potential by decreasing the binding of PrP^C^ to PINK1. **A.** Immunoprecipitates with anti-PINK1 were analyzed treatment Healthy-hMSCs with or without si-PRNP and CKD-hMSCs by western blot using an antibody that recognized PrP^C^. **B.** The positive of MitoSOX was counted by FACS analysis of staining treatment Healthy- hMSCs with or without si-PRNP and CKD-hMSC. **C, D.** Complex I and complex IV activities were analyzed after treatment of Healthy-hMSCs with or without si-PRNP and in CKD-hMSCs by ELISA. Values represent the mean ± SEM. \**p* < 0.05 vs. Healthy-hMSC.

**Figure EV3.**
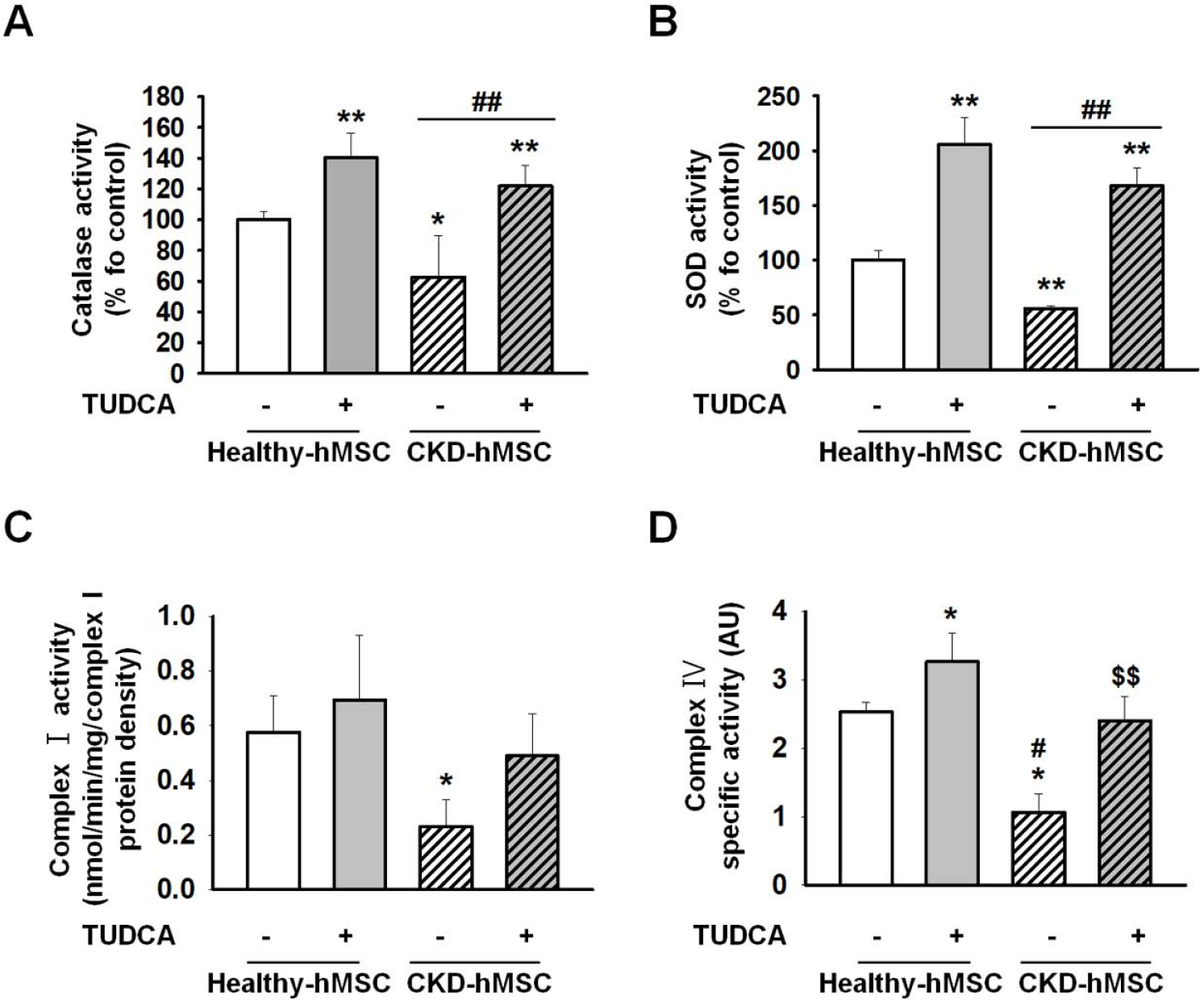
Treatment of Healthy-hMSCs with si-PRNP decreased mitochondrial membrane potential by decreasing the binding of PrP^C^ to PINK1. **A, B.** Catalase and SOD activity were assessed treatment Healthy-hMSCs and CKD-hMSCs with or without TUDCA. Values represent the mean ± SEM. \**p* < 0.05, and \**p* < 0.01 vs. Healthy-hMSC. ##*p* < 0.01 vs. TUDCA-treated Healthy-hMSCs. $$*p* < 0.05 vs. CKD-hMSC. **C, D.** Complex I and complex IV activities were analyzed treatment Healthy-hMSCs and CKD-hMSCs with or without TUDCA by ELISA. Values represent the mean ± SEM. \**p* < 0.05 vs. Healthy-hMSC. #*p* < 0.05 vs. TUDCA-treated Healthy-hMSC. $$*p* < 0.05 vs. CKD-hMSC

